# mTOR Inhibition Suppresses Salinomycin-Induced Ferroptosis in Breast Cancer Stem Cells by Ironing Out Mitochondrial Dysfunctions

**DOI:** 10.1101/2023.05.02.539040

**Authors:** Emma Cosialls, Emeline Pacreau, Sara Ceccacci, Rima Elhage, Clémence Duruel, Christophe Desterke, Kevin Roger, Chiara Guerrera, Romane Ducloux, Sylvie Souquere, Gérard Pierron, Ivan Nemazanyy, Mairead Kelly, Elise Dalmas, Yunhua Chang, Vincent Goffin, Maryam Mehrpour, Ahmed Hamaï

## Abstract

Ferroptosis constitutes a promising therapeutic strategy against cancer by efficiently targeting the highly tumorigenic and treatment-resistant cancer stem cells (CSCs). We previously showed that the lysosomal iron-targeting drug Salinomycin (Sal) was able to eliminate CSCs by triggering ferroptosis. Here, in a well-established breast CSCs model (human mammary epithelial HMLER CD24^low^/CD44^high^), we identified that pharmacological inhibition of mechanistic target of rapamycin (mTOR), suppresses Sal-induced ferroptosis. Mechanistically, mTOR inhibition modulates iron cellular flux and prevents the iron and ROS bursts induced by Sal. Besides, integration of multi-omics data identified mitochondria as a key target of Sal action. We demonstrated that mTOR inhibition prevents Sal-induced mitochondrial functional and structural alteration, and that Sal-induced metabolic plasticity is mainly dependent on the mTOR pathway. Overall, our findings provide experimental evidences on the detailed mechanisms of mTOR as a crucial effector of Sal-induced ferroptosis, and gives proof-of-concept that careful evaluation of such combination therapy (here mTOR and ferroptosis co-targeting) is required for effective treatment.

**GRAPHICAL ABSTRACT:** 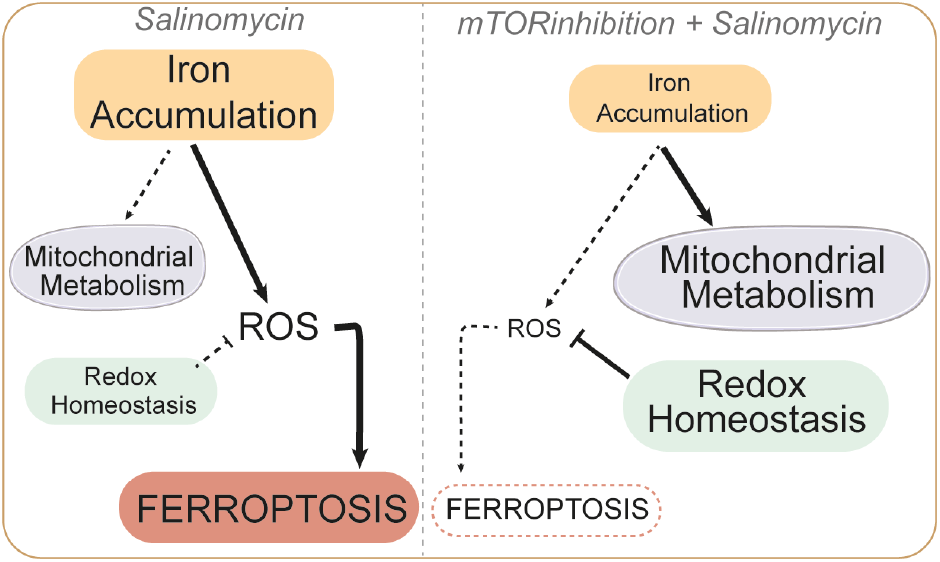

## INTRODUCTION

Tumor relapse and metastasis, along with increased resistance to conventional therapies, are a major clinical challenge in curing breast cancer. The therapeutic failure is thought to be caused by a sub-population of highly tumorigenic cells with stem cell properties, termed cancer stem cells (CSCs) ^1–3^. We previously showed that breast CSCs are highly sensitive to ferroptosis, a non-apoptotic and iron-dependent cell death, while being resistant to conventional therapy compared to bulk tumor cells ^4,5^. Salinomycin (Sal), a polyether antibiotic, selectively kills breast CSCs by ferroptosis *in vitro* and *in vivo.* Mechanistically, we have shown in well-established breast CSCs model (human mammary epithelial HMLER CD24^low^/CD44^high^)^6,7^ that Salinomycin triggers ferroptosis by sequestering iron in the lysosomes^4,8^. However, since the specific regulatory mechanisms of Sal-induced ferroptosis are still unexplored.

Recently, accumulating evidence has identified the mTOR pathway, as a critical regulator of ferroptosis^9^, sometimes negative^10–12^, and other times positive^13,14^. This oncogenic pathway is one of the most frequently dysregulated pathways in cancer. It is a master controller of cell growth, survival, and metabolism, and is activated by several factors including growth factors and nutrients.

The present study confirms the crucial role of mTOR in ferroptosis, and indicates that mTOR inhibition suppresses the therapeutic effect of Sal in breast CSCs. Mechanistically, inhibition of mTOR prevents the Sal-induced iron burst and thereby limits iron-mediated oxidative stress. Furthermore, an integrated metabolomics and proteomics approach provides new insights into mitochondria as a key target of Sal action, leading to profound alterations in mitochondrial metabolic pathways prevented by mTOR inhibition. Overall, our work supports that Sal-induced metabolic plasticity is mainly dependent on the mTOR pathway, and that its inhibition exerts a protective role against ferroptosis by modulating iron homeostasis, preventing the Sal-induced metabolic burden while decreasing the accumulation of damaged mitochondria. Furthermore, our study underlines that metabolic reprogramming regulates ferroptosis sensitivity, thus opening new opportunities to treat tumors unresponsive to therapies.

## RESULTS

### mTOR Inhibition Protects Cells from Sal-Induced Cell Death

To investigate if mTOR impacts Sal-induced ferroptosis, HMLER CD24^low^ cells were treated with the ATP-competitive mTOR inhibitor Torin^15^ in combination with Sal during 96h. Intriguingly, Torin treatment potently suppressed Sal-induced cell death (**Figure 1A and 1B**). Torin blocked as expected the phosphorylation of the mTORC1 substrates: S6, p70S6K and 4EBP1 (**Figure 1C-quantifications in S1A**). Of note, Sal treatment alone activated the phosphorylation of S6 and S6K (**Figures 1C-quantification in S1A**). To confirm the protective effect of mTOR inhibition, other inhibitors were tested: ATP-competitive inhibitors (Torin-2 or AZD8055) also inhibited Sal-induced cell death to the same extent as Torin, while the well-known mTORC1 inhibitor Rapamycin had a weaker effect (**Figures S1B and S1C**). Next, siRNA targeting subunits of mTORC1 (si*RAPTOR*, regulatory-associated protein of mTOR) or mTORC2 (si*SIN1*, mammalian stress-activated protein kinase-interacting protein 1) were used (**Figure 1D**). Upon Sal treatment, only cells knocked-down for RAPTOR prevented Sal-induced cell death (**Figure 1E**). Hence, these data suggest that the suppression of Sal-induced cell death is driven specifically by mTOR inhibition, mainly through mTORC1. Furthermore, the use of MHY1485, a compound described to activate mTOR ^16^, potentiated Sal-induced cell death (**Figure 1F**). Overall, these data highlight the crucial role of mTOR signaling in the regulation of Salinomycin-induced cell death.

**Figure 1.**
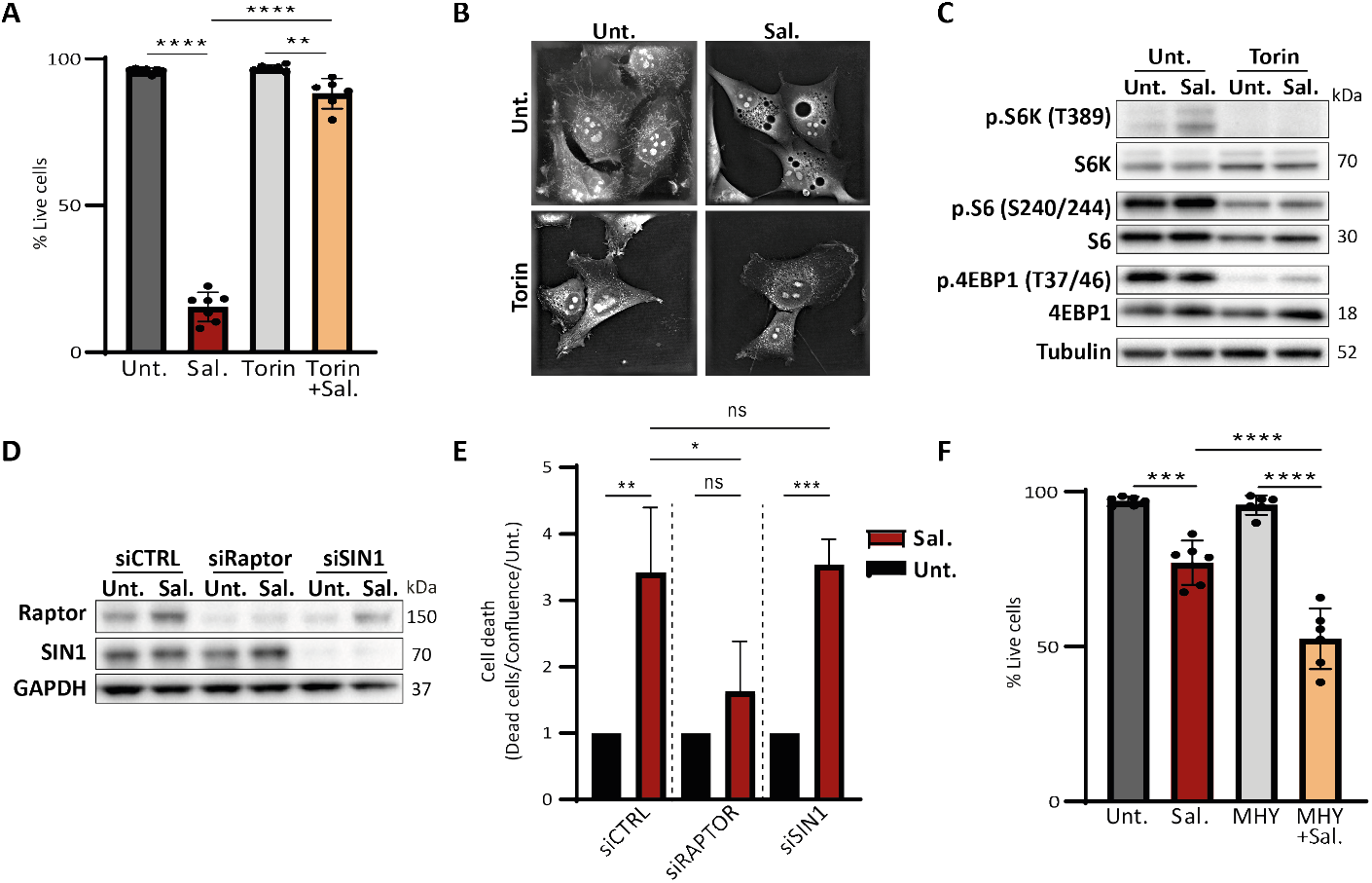
mTOR inhibition prevents cell death induced by Sal. **(A-C)** HMLER CD24L cells were treated with Sal (500 nM), Torin (250 nM) or combination of both. **(A)** Cell death determined by dapi staining coupled with flow cytometry (FC) after 96h (n=7). **(B)** Live cell imaging by 3D holotomographic microscopy (Nanolive) after 48h of treatment. **(C)** Immunoblotting for the indicated mTOR-related protein after 48h. Tubulin used as loading control. **(D-E)** HMLER CD24L knockdown for either *RAPTOR* or *SIN1* and then treated as indicated. **(D)** Immunoblotting for the indicated silenced protein after 48h. GAPDH used as loading control. **(E)** Acquisition of cell viability using Incucyte Live-cell analysis, dead cells were counted by Incucyte® Cytotox Dye Probe over cell confluence normalized on each untreated condition after 120h of treatment (n=3). **(F)** HMLER CD24L were treated with either Sal (500 nM), MHY1485 (5 μM) or combination of both for 72h and cell death was determined by dapi staining coupled with FC (n=7). Data are presented: mean ±SD, ANOVA test: ****p<0.0001; ****p<0.002; ****p<0.001

### mTOR Inhibition Prevents ROS and Iron Accumulation

**Figure 2A** summarizes iron homeostasis and the mechanisms by which Sal leads to iron accumulation, excessive ROS production and lipid peroxidation, and ultimately triggers ferroptosis^17,4,18^. We therefore investigated the effect of mTOR inhibition on these features after 48h of treatment. Firstly, Sal treatment markedly increased ROS levels, including both global ROS and hydroxyl radicals, and iron level, as expected; while co-treatment with Torin significantly reduced ROS and iron levels (**Figures 2B, 2C and 2D**). However, lipid peroxidation level was not significantly impacted by treatments (**Figure S2A**).

**Figure 2.**
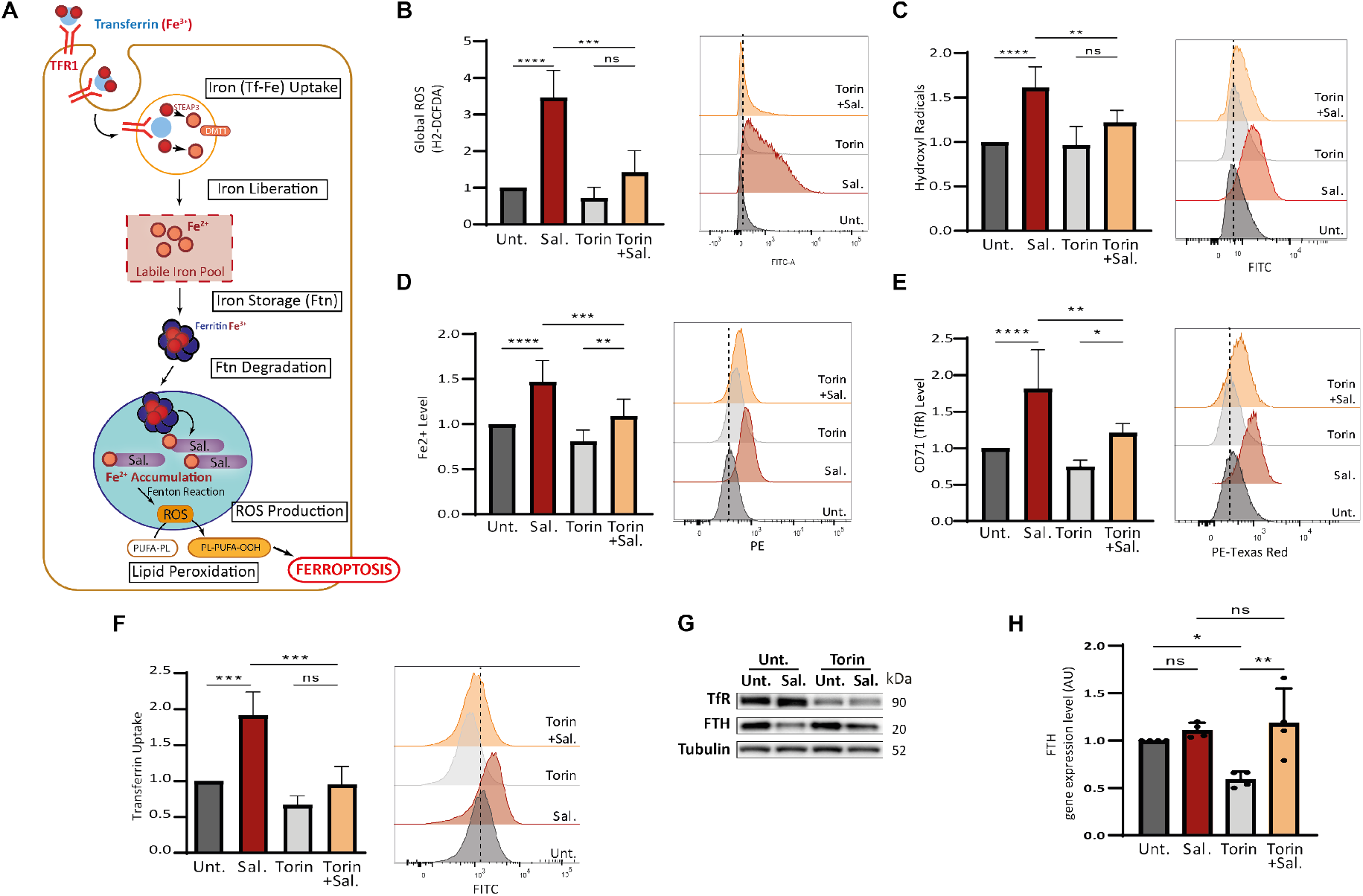
mTOR inhibition reduced pro-ferroptotic hallmarks induced by Sal. **(A)** Iron entry into the cell is mediated by the binding of Fe^3+^-Tf complex to TFR and its subsequent endocytosis. Iron is released under the acidic environment of the endosome, and is reduced to Fe2+ by STEAP3 and transported into the cytosol by DMT1. Free iron, constituting the labile iron pool, can be used (in mitochondria, …), exported or stored in ferritin. Low iron availability triggered the degradation of ferritin in mitochondria. Salinomycin sequestered iron in the lysosomes, triggering a cellular iron-depletion response that induced an increase iron entry, and an increase ferritin degradation, it results in an accumulation of iron in the lysosome and a subsequent massive ROS production (by the Fenton reaction) leading to a considerable lipid peroxidation and ultimately cell death. **(B-G)** HMLER CD24L were treated with Sal, Torin or combination of both for 48h. **(B)** Global ROS level determined by H2-DCFDA staining coupled with FC (n=4). **(C)** Hydroxyl Radical level determined by HPF staining coupled with FC (n=6). **(D)** Fe^2+^ level determined by FerroOrange staining coupled with FC (n=9). **(E)** TFR level determined by anti-CD71 staining coupled with FC (n=7). **(F)** Transferrin Uptake determined by Alexa-488-Transferin staining coupled with FC (n=4). **(G)** Immunoblotting for the indicated iron-related protein. Tubulin used as loading control. **(H)** FTH gene expression level by RT-qPCR. Data are presented: mean ±SD, ANOVA test: ****p<0.0001; ****p<0.002; ****p<0.001

To determine the mechanisms by which Torin protects against Sal-induced iron overload, iron entry was first considered in cells treated during 48h. Confirming previous observations^4^, Sal elicited iron uptake via increased transferrin (Tf)-Uptake and CD71 expression at the cell surface (**Figures 2E and 2F**) as well as at the total protein (**Figure 2G-quantifications in S2B**) and mRNA (**Figure S2C**) levels. In contrast, Torin alone, or in combination with Sal, prevented iron uptake (**Figures 2E, 2F and 2G**). Cellular iron content also results from the degradation of ferritin, an iron storage molecule, which is recycled/degraded in lysosome under low iron levels, namely ferritinophagy ^19^ (**Figure 2A**). Consistent with our previous report ^4^, Sal triggered ferritinophagy as shown by decreased FTH protein level (**Figures 2G – quantification in S2D**) and increased FTH mRNA level (**Figure 2H**). Remarkably, although Torin is a strong inducer of autophagy and inhibitor of protein synthesis, co-treatment with Torin upregulated the level of FTH protein (but not its mRNA) compared to Sal alone (**Figures 2G and 2H – quantification in S2D**). To investigate further ferritin degradation, cells were additionally treated with a specific autophagy inhibitor (Hydroxychloroquine, HCQ)^20^, as confirmed by the accumulation of p62 and LC3 proteins (**Figure S2E**). HCQ and Sal co-treatment induced an increase in FTH protein level compared to Sal alone. However, it did not affect Sal-induced cell death (**Figures S2E and S2F**), suggesting that blocking FTH degradation is not sufficient to prevent cell death. Co-treatment of HCQ with Torin (with or without Sal) did not increase the FTH protein level compared to Torin (with or without Sal), indicating that mTOR inhibition *per se* prevents the proteostasis of FTH (**Figure S2E**). Overall, these data indicate that mTOR inhibition prevent ROS production and iron homeostasis dysregulation induced by Salinomycin.

### Salinomycin Drastically Disrupts the Expression of Mitochondrial Proteins which is Protected by mTOR Inhibition

To further investigate the key actors involved in Sal action and also in ferroptosis suppression by mTOR inhibition, we carried out a proteomic analysis by using mass spectrometry on cells treated for 48h. In total, we identified and quantified 6,396 proteins. Among them, the level of 1,919 proteins were found to be significantly different after 1-way ANOVA analysis (FDR 1% and s0=1) between the four conditions, and an unsupervised hierarchical clustering analysis identified 6 main clusters. Cluster A showed proteins downregulated by Sal treatment compared to control, and upregulated by co-treatment with Torin compared to Sal alone (**Figure 3A**). Functional annotation of this cluster using molecular function (MF), cellular component (CC) and biological process (BP) terms pointed out to mitochondrial function and ribosomal activity (**Figures 3B and S3**).

**Figure 3.**
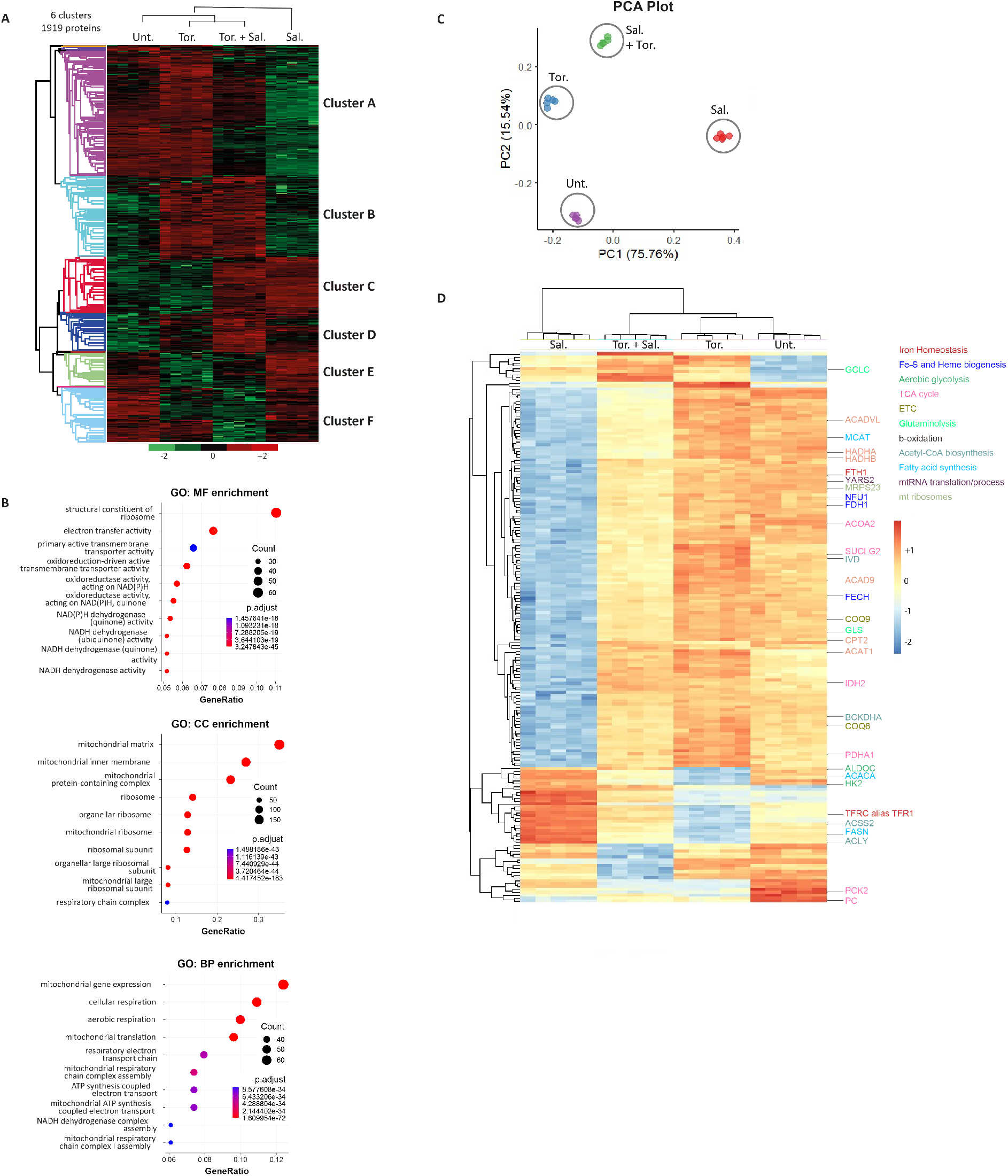
Salinomycin drastically impairs the expression of mitochondrial proteins which is protected by mTOR inhibition. HMLER CD24L cells were treated or not with Sal (500 nM) or/and Tor. (250 nM) for 48h. **(A)** Unsupervised clustering performed with the 1919 proteins differently modulated between the different conditions of treatment from proteomic data (ANOVA test, FDR<0.01, S0=1). 6 main distinct clusters have been identified. Each column is a sample; each row a protein. The color scale indicates the protein expression value (green: lowest; red: highest). The intensity of each protein corresponds to the relative abundance of individual proteins by liquid chromatography-mass spectrometry (nanoLC-MS/MS). Proteins were clustered using Perseus software with Euclidean distances. (n=5 replicates). **(B)** Dot plot of the best Gene Ontology (top 10 terms regarding best adjusted p-value) for Molecular Function (MF), for Cellular Component (CC), and for Biological Process (BP) obtained from the differentially expressed proteins of cluster A. The size of the circles represents the number of proteins (Count) found enriched for each corresponding term. **(C-D) Identification of a 187-protein-based discriminant signature of the four experimental conditions investigated by proteomic analysis. (C)** Principal component analysis performed with the 187-protein based signature supervised by machine learning. **(D)** Unsupervised clustering performed with the 187-protein based signature.

To determine the key pathways involved in ferroptosis protection through mTOR inhibition, a minimal proteomic signature of 187 proteins was identified by processing a supervised machine learning on the 1,919 proteins differentially expressed. All the steps of the analysis are detailed in Materials and Methods and illustrated in **Figure S4A, B and C**. By unsupervised principal component analysis, the 187-proteins signature successfully discriminated the samples on the first two principal components accounting for more than 91% of the signature variance (**Figure 3C**). By unsupervised clustering, the expression of the 187 proteins discriminated well the samples groups (**Figure 3D**). Most of the proteins contained in this signature are downregulated by Sal treatment (150/187) as compared to control, and of note, their levels are upregulated upon co-treatment with Torin (**Figure 3D**). Among the signature, we found proteins involved in iron homeostasis, including FTH and TfR, which are oppositely regulated by Sal alone versus co-treatment with Torin, (**Figure 3D** – proteins written in dark red) in agreement with our previous results. Furthermore, proteins involved in mitochondrial iron metabolism and essential for mitochondrial functions, including Fe-S or heme biogenesis, were found to be downregulated by Sal, and in comparison, up-regulated by co-treatment (**Figure 3D** – proteins written in blue). Once again, this data highlights that the iron dysregulation induced by Sal is attenuated by co-treatment with Torin. Besides, functional enrichment analysis performed with KEGG database on the network reported a strong enrichment in metabolic pathways associated with mitochondrial pathways (**Figures S4D and E**). In line with this, we also found in this signature many proteins involved in mitochondrial - pathways or -related pathways (e.g., TCA cycle, electron transport chain, glycolysis and glutaminolysis, β-oxidation) and in mitochondrial biogenesis (e.g., mtRNA translation/process and in mt ribosomes) as highlighted in **Figure S5**. Considering the importance of iron metabolism in mitochondrial function, this suggests that in our model iron dysregulation drives mitochondrial alterations and triggers ferroptosis.

### Integration of -Omics Data Highlighted a Metabolic Shift under Sal treatment, prevented by mTOR inhibition

Ferroptosis has been described as a metabolism-associated cell death^21^, and given the observed overall impact seen on mitochondrial function, we postulated that the protective effect of mTOR inhibition could be mediated by changes in cellular metabolism. To this end, a targeted analysis of individual metabolites using liquid chromatography–mass spectrometry (LC-MS) was performed on cells treated for 48h. In order to identify the key pathways involved in ferroptosis execution by Sal and those that lead to its blockage by Torin, we decided to integrate metabolomics and proteomics data (**Figure 4**). **Figures 4A and B** show the heatmap of proteins and metabolites level involved in mitochondrial pathways (OXPHOS/TCA) and mitochondrial-related pathways (glycolysis and glutaminolysis) in cells treated for 48h. Consistent with the decreased level of TCA- and ETC- related enzymes, Sal treatment dramatically reduced the level of the majority of TCA cycle intermediates with an increase in acetyl-Coenzyme A (acetyl-CoA) indicative of impaired TCA activity (**Figure 4A-**proteins in pink **and 4B**). In addition, the accumulation of succinate shown by the increase in succinate/fumarate ratio and the impairment of Succinate Dehydrogenase SDH (or complex II) level, supports this TCA- and ETC- alteration mediated by Sal (**Figure 4B,** *lower panel*). Therefore, as summarized in **Figure 4C**, Sal treatment inhibits both TCA and ETC pathway, yet the addition of Torin seems to restore both.

**Figure 4.**
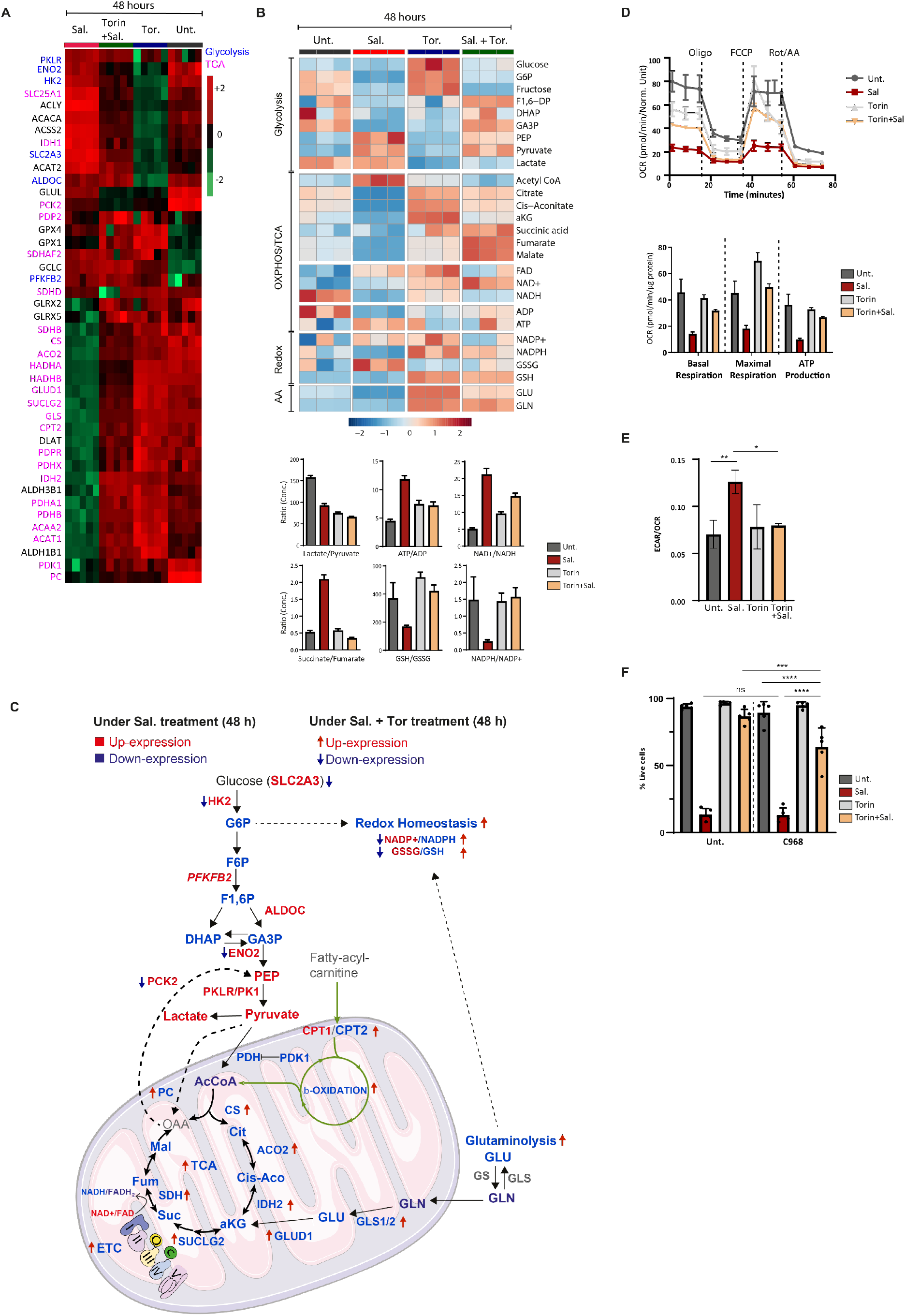
mTOR inhibition interferes with sal-mediated enhanced aerobic glycolysis and restores the mitochondrial-associated metabolic pathways. **(A)** Heatmap comparing relative levels of proteins related to TCA cycle (written in pink) and glycolysis (written in blue) in HMLER CD24L cells in response to Sal, or Tor. alone, and combination treatments for 48 h. Color key indicates protein expression value (green: lowest; red: highest). Proteins were clustered using Perseus software. **(B)** *Upper:* Heatmap comparing relative levels of metabolites related to OXPHOS or TCA cycle and to glycolysis and also to glutaminolysis in HMLER cells in response to Sal, or Tor. alone, and combination treatments for 48 h. Color key indicates metabolite expression value (blue: lowest; red: highest). Metabolites were clustered using Metaboanalyst software. *Lower:* Graphs represent the mean of lactate/pyruvate, ATP/ADP, NAD+/NADH, Succinate/Fumarate ratios indicating the use of glycolysis or OXPHOS. GSH/GSSG and NADPH/NADP+ ratios are indicators of oxidative stress. **(C)** Summary of the differential level of metabolites and proteins associated with the OXPHOS/TCA cycle, with glycolysis, with b-oxidation, with glutaminolysis and redox homeostasis under Sal treatment (terms) and under combination treatment (arrows). Blue and red colors indicate the downregulated and upregulated level, respectively. **(D)** Seahorse-based measurements of OCR (*upper panel*) in HMLER CD24L cells incubated under treatments as indicated for 48 h, normalized to total protein levels. ***Upper panel:*** Oligomycin, carbonyl-cyanide-4-(trifluoromethoxy)phenylhydrazone (FCCP), and antimycin A were serially injected to measure ATP production, maximal respiration, and basal respiration, respectively, as indicated in the graph. **(E)** Basal ECAR/OCR ratio in HMLER CD24L cells treated as indicated for 48 h. **(F)** HMLER CD24L treated during 96h with either Sal (500 nM), Torin (250 nM) or combination of both in presence or in absence of glutaminolysis inhibitor (C968 (20 DD)). Cell death determined by dapi staining and flow cytometry. n=3 independent experiments (with at least duplicate) Data are presented: mean ±SD, ANOVA test: ****p<0.0001; ****p<0.002; ****p<0.001

Next, regarding glycolysis, we found that Sal treatment (compared to untreated) decreased the lactate-to-pyruvate ratio (anaerobic glycolysis marker) and increased ATP/ADP ratio (**Figure 4B,** *lower panel*), as well as upregulated cytoplasmic glycolytic enzymes (**Figure 4A** - proteins in blue). Whereas the pyruvate dehydrogenase complex (PDH) that converts pyruvate into Acetyl-CoA for TCA cycle is downregulated (**Figure 4B,** *lower panel*). To go further, we assessed mitochondrial and glycolytic activity by measuring the oxygen consumption rate (OCR) and the extracellular acidification rate (ECAR) respectively using Seahorse-based assays. (**Figure 4D**). We confirmed Sal-induced ETC dysregulation with the significant alteration of both basal and maximal respiration and ATP production, all prevented by co-treatment with Torin (**Figure 4D**). Besides, the basal ECAR-to-OCR ratio increased under Sal treatment demonstrating higher glycolytic to mitochondrial activity, which was reversed under Torin treatment (**Figure 4E**). In short, Sal-treated cells shift to a higher aerobic glycolysis to produced ATP, while co-treated cells restored the expression of the TCA-associated enzymes, ETC-related proteins and ATP-production by OXPHOS (**Figure 4C**).

### Activation of Glutaminolysis Seems to Drive Ferroptosis Suppression by mTOR Inhibition

Apart from glycolysis, TCA can also be sustained by glutamine anaplerosis (**Figure 4C**). Along with the decrease in TCA activity, Sal treatment also reduced the level of glutaminolysis-related metabolites and enzymes (GLS, GDH and GLUD1) (**Figure 4A and B**). In addition, glutamine is also involved in redox-homeostasis through the production of GSH and NADPH, two essential substrates for the antioxidant defenses. Sal treatment resulted in a decrease in the NADPH-to-NADP+ ratio and a decrease in reduced-to-oxidized glutathione ratio (GSH/GSSG) (**Figure 4B,** *lower panel*). Importantly, all of these modulations were restored upon co-treatment with Torin, suggesting increased glutaminolysis and improved management of oxidative stress, consistent with our previous results. To determine whether activation of glutaminolysis contributes to Torin protection against ferroptosis, cells were additionally treated with the glutaminase (GLS) inhibitor: Compound C968 (C968). Co-treatment with C968 did not affect Sal-induced cell death (**Figure 4F**). However, it sensitized cells to Sal-induced cell death upon co-treatment with Torin. Therefore, inhibition of glutamine anaplerosis prevents Torin treatment from suppressing Sal-induced cell death. As summarized in **Figure 4C**, Sal-treated cells undergo a profound metabolic reprogramming which could result in lower redox homeostasis. On the contrary, upon mTOR inhibition cells showed a greater oxidative metabolism, with increased TCA- and ETC- activity, as well as overactivation of glutaminolysis that contribute to the Torin-induced protection against ferroptosis.

### mTOR Inhibition Overcomes Salinomycin-induced Mitochondrial Dysfunction

Given that Sal-treatment massively affects mitochondria-related processes, we sought to better understand how the different treatments impinge on mitochondria (**Figure 4C**). Similarly, with respect to mitochondrial respiration, the treatment decreased the protein level of the complexes I, II, III and IV of the ETC, whereas co-treatment with Torin prevented this reduction (**Figures 5A, S6A - quantification in S6B**). To further investigate, mitochondrial membrane potential (ΔΨ) (generated by complexes I, III, and IV) was measured using mitochondrial probes: MitoCMXRos accumulates in negatively charged mitochondria and MitroTracker accumulates in the matrix. By analyzing the ratio of MitoCMXRos-to-MitoTracker, we found that Sal treatment increased ΔΨ, while co-treatment with Torin reduced it (**Figure 5B**). Interestingly, mitochondrial mass was not affected by treatments (**Figure 5B,** *right panel*). Defective mitochondria are more likely to produce ROS that could promote lipid peroxidation. Therefore, the lipid-ROS level was measured using a fluorescence mitochondria-targeted lipid peroxidation probe (MitoPerOx) ^22^. Sal treatment alone increased Mt lipid peroxidation, while co-treatment with Torin reduced it (**Figure 5C**). Overall, these data indicate that mTOR inhibition prevents Sal-mediated mitochondrial dysfunction and oxidative stress.

**Figure 5.**
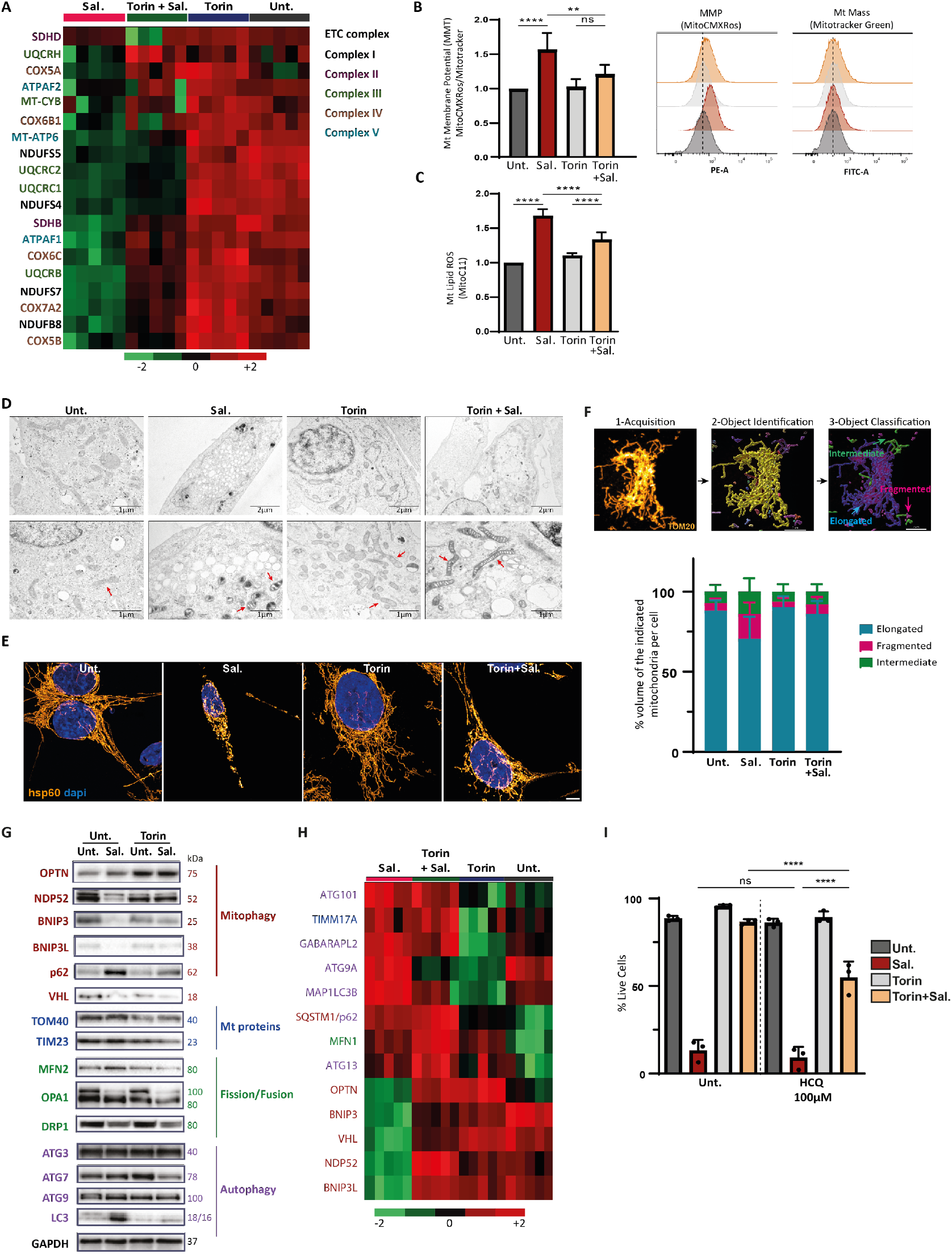
mTOR inhibition prevents mitochondrial damages induces by Sal. HMLER CD24L were treated with either Sal, Torin or combination of both for 48h. **(A)** Heatmap of ETC proteins from proteomics data. Color key indicates protein expression value (green: lowest; red: highest). **(B)** Mitochondrial membrane potential measured with MitoCMXRos and Mitotracker Green staining. Data are shown as a ratio of MitoCMXRos over Mitotracker Green (n=5). **(C)** Mitochondrial lipid peroxidation was measured by Mito-C11 probe staining couple with FC. Representative histogram of oxidized-Mito-C11 (FITC channel) intensity level in cells (n=3). **(D)** TEM images, red arrows show mitochondria. **(E)** Immuno-staining of mitochondria with hsp60 and images acquired using Confocal Leica TCS SP5. Objective: 63x. Scale bar:5μm. **(F)** *Upper panel,* steps of the workflow developed to analysis mitochondrial network: first, a machine learning (ML) identified mitochondria as object, then a second ML classified mitochondria into three classes depending on their shapes: fragmented, intermediary and hyperfused. *Lower panel,* Quantification of mitochondrial network analysis, data shown as % area of each class for each cell. Analysis on >25 cells from 2 independent experiments. **(G)** Immunoblotting for the indicated autophagic/mitophagic proteins and Mt/dynamic proteins. GAPDH was used as the loading control. **(H)** Heatmap of autophagic/mitophagic proteins and Mt/dynamic proteins from proteomics data. Color key indicates metabolite expression value (green: lowest; red: highest). Protein terms and functions indicated with the same color are corresponding for (**G**) and (**H**). (**I**) HMLER CD24L cells were treated with Sal or/and Tor. in presence or in absence of HCQ (100 μM) for 144 h. Cell death determined by dapi staining coupled with flow cytometry (FC). The graph represents the mean (± SEM) of three independent experiments. One-way ANOVA test. *p < 0.05; **p < 0.01; ***p < 0.001; ****p < 0.0001. ns, not significant.

### Salinomycin Profoundly Alters Mitochondrial Network while mTOR Inhibition Restores a Reticular Network

Given that mitochondrial activity is altered by Sal treatment, we qualitatively evaluated the mitochondrial network during treatment. Firstly, an ultrastructure analysis of mitochondria by transmission electron microscopy (TEM) revealed that Sal- treated cells had smaller mitochondria with profound alteration in their structural integrity such as darker matrix and irregular cristae compared to untreated cells (**Figure 5D -** red arrows show mitochondria). On the contrary, cells treated with Torin showed much less affected mitochondria, with an enlargement of mitochondrial cristae. A mitochondrial network analysis by immunostaining with a mitochondrial matrix protein (hsp60, Heatshock protein 60) confirmed that Sal treatment dramatically affected the mitochondrial network with fragmented mitochondria, while a phenotype close to untreated cells was observed in co-treated cells (**Figure 5E**). To take this step further, a workflow for the analysis of 3-dimensional mitochondrial networks was processed as precisely decribed in the Materials and Methods section. **Figure 5F** *upper panel,* briefly details the steps of the workflow. This analysis highlighted the increased proportion of fragmented mitochondria in Sal-treated cells compared to untreated, Torin- or co-treated cells (**Figures 5F,** *lower panel* **and S6C**). Overall, these data indicate that mTOR inhibition prevents the profound alteration in the mitochondrial morphology induced by Salinomycin.

### mTOR Inhibition Prevents Mitochondrial Dynamic Alteration induced by Salinomycin

The maintenance of the mitochondrial network is a highly dynamic process regulated by fusion and fission events, as well as the elimination of damaged mitochondria, a process called mitophagy ^23^. Since Torin is known to activate mitophagy^24^, and given the impact of treatment on the mitochondrial network, we decided to study key actors involved in mitochondrial dynamics. Proteomic data as well as western blot analysis showed downregulation of several mitophagy proteins (including BNIP3/3L, OPTN, NDP52) in Sal-treated cells, whereas Torin treatment restored it (**Figures 5G and 5H**). Besides, as described in our previous work^25^, although ATG initiator proteins were upregulated, an accumulation of LC3-II and p62 was observed following Sal treatment indicating inhibition of autophagic flux that is restored upon co-treatment. Then, mitochondrial DNA (mtDNA), cleared during mitophagy to avoid its accumulation, was measured by qPCR^26^. Co-treatment decreased the mitochondrial mtDNA mass as compared to Sal alone (**Figure S6D**) supporting the activation of mitophagy in co-treated cells.

In line with these results, TEM images acquisition revealed that cells co-treated with Torin exhibited advanced autophagic degradative vacuoles/autolysosomes containing damaged organelles, including a structure with the same dark density as the damaged mitochondria (**Figure S6E**). The importance of Torin-induced clearance of damaged mitochondria in the protection against ferroptosis was examined by further treating cells with HCQ (100 DM) to block mitophagy ^27 28^. HCQ efficiently inhibits the late stage, i.e. the fusion of mitophagosomes with lysosomes and blocks the degradation of mitochondria. HCQ treatment after 144h significantly decreased the viability of co-treated cells, restoring their sensitivity to Sal-induced cell death (**Figure 5I**). Of note, HCQ alone or with Torin did not affect the Sal-induced ferroptosis. Taken together, these results suggest that the autophagic removal of Sal-induced damaged mitochondria contributes to the restoration of mitochondrial functions and to the ferroptosis protection induced by mTOR inhibition.

## DISCUSSION

In the present study, we examined the key role of mTOR pathway in ferroptosis induced by Salinomycin. In particular, mTOR inhibition interferes with the Sal-induced iron dysregulation and metabolic rewiring, thereby decreasing the sensitivity of CSC to ferroptosis, and notably preserving the integrity of the mitochondrial network and promoting the clearance of damaged mitochondria (**Figure 6**).

**Figure 6.**
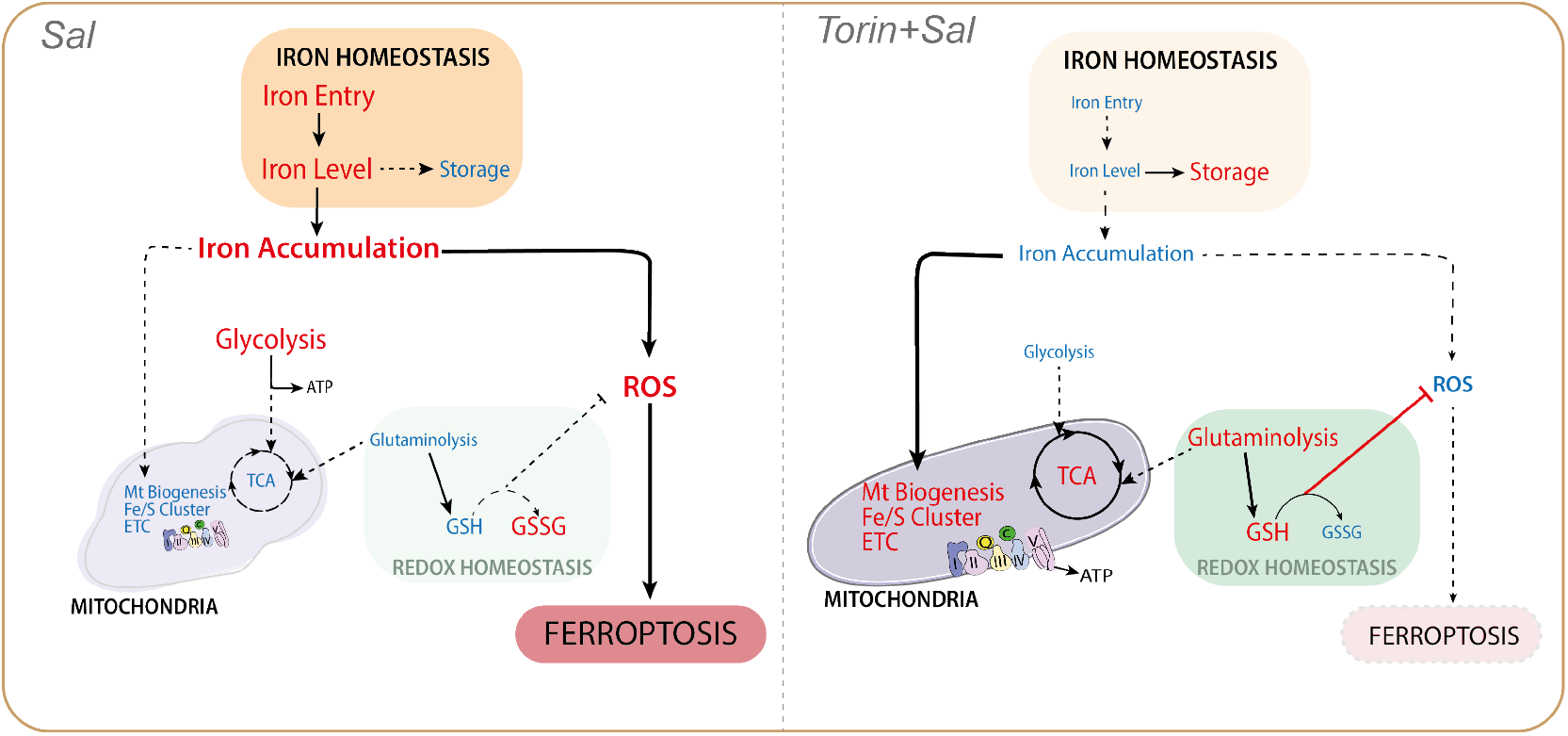
Schematic Proposal Model of the Suppression of Ferroptosis by mTOR Inhibition. **(A)** Pathways regulated upon Sal treatment compared to control. Mechanistically, Sal induces a burst of iron and ROS, and triggers a metabolic shift by decreasing the level of mitochondrial proteins and the activity of mitochondrial metabolic pathways. Besides, Sal inhibits redox homeostasis, which leads to even more ROS production, and subsequently triggers ferroptosis. **(B)** Pathways regulated upon co-treatment with Torin and Sal compared to Sal alone. In contrast, inhibition of mTOR prevents the Sal-induced accumulation of iron and ROS, as well as the functional and structural alteration of mitochondria. Besides it activates glutaminolysis to improve redox defenses which leads ultimately to the inhibition of ferroptosis. Blue and red colors indicate the downregulated and upregulated level, respectively.

The accumulation of iron within cells through Sal-induced increased iron entry is impaired by mTOR inhibition, consistent with the work of Bayeva et al.^29^ showing that mTOR inhibition decreases the stability of TfR1 mRNA and alters cellular iron flux. This suggests that less Fe^2+^ are mobilized for Fenton chemistry, consistent with the decreased ROS level. Overall, these data indicate that mTOR inhibition affects various molecular regulators related to iron homeostasis. Metabolomic and proteomic analyses highlighted mitochondria as a key regulator of Sal-induced ferroptosis supporting the numerous studies showing that mitochondria are a major hub of ferroptosis^30–32^. Interestingly, Smethurst DG. et al., recently identified that iron- promotes ribosomes RNA degradation under oxidative stress in yeast^33^. Besides, ROS accumulation impairs the mitochondria integrity including mtDNA maintenance and mtRNA processing^34^, and the inhibition of Sal-induced accumulation of mtROS under mTOR inhibition, may limit the severe mitochondrial damages, including by increasing de novo GSH synthesis ^13^.

Furthermore, some recent data showed that treatment with a Sal-derivative termed AM5, leads to a decrease in mitochondrial iron level in acute myeloid leukemia ^35^. Interestingly, the expression of Frataxin (FXN), a protein involved in Fe/S cluster biogenesis, has been recently identified as a negative regulator of ferroptosis, and is upregulated upon mTOR inhibition^36^. The effect of Sal on mitochondrial dysfunction is likely mediated by some downregulated Fe/S proteins, that are upregulated with co-treatment. Particularly, NEET proteins including both CISD1 (also termed mitoNEET) and CISD3 (also known as Mine2/MiNT) (which regulate iron and ROS homeostasis) have been demonstrated to protect cells from mitochondrial damage in ferroptosis^37,38^. Among Fe/S-containing enzymes, loss of Succinate Dehydrogenase (SHDB/C) (protein related to TCA/ETC), has been shown to affect iron homeostasis and promote ferroptosis^39^. Thus, Salinomycin drastically induces a profound metabolic reprogramming toward aerobic glycolysis that is characterized by the downregulation of mitochondrial metabolic pathways related to TCA cycle/ETC activity. In agreement with previous report^40^, inhibition of mTOR could also restore TCA cycle activity by anaplerosis, notably via glutaminolysis, which could also be involved in the maintenance of redox homeostasis through *de novo* GSH synthesis, as described above.

We revealed by machine learning analysis of the mitochondrial network that Sal increases intermediate and fragmented mitochondria while cells treated with mTOR inhibitor display a tubular mitochondrial network. However, the dysregulation of mitochondrial dynamics proteins induced by Sal treatment does not *fit* with the observed fragmented networks, which raises the question: by which mechanisms? A possible explanation is the impact of redox sensible post-translational modifications on their activity. For example, it has been shown that only the mature L-isoform (L∼100kDa vs S∼80kDa) of the dynamin-like GTPase OPA1 has a mitochondrial fusion stimulating activity ^41^. Or again, that under oxidative stress S-nytrosylation increases the GTPase activity of DRP1 ^42^. Besides, mitochondrial membrane hyperpolarization upon Sal treatment may contribute to enhanced mitochondrial fission. In line with this, we found that the inhibition of mitophagy contributes to restore cell sensitivity to Sal-cell death upon mTOR inhibition. Interestingly, NDP52 has been recently identified as a redox sensor in autophagic clearance of damaged mitochondria^43^. Our results show that Sal treatment dramatically affects mitochondrial dynamic and function, and might overcomes the cell’s capacity to cleared damaged mitochondria even under a massive oxidative stress, mechanisms which are, in several ways, prevented by mTOR inhibition. These data are in agreement with other studies showing that mTOR inhibition protects against mitochondrial diseases through mitophagy activation ^44–46^.

As mTOR inhibitor in monotherapy has efficacy in several types of cancer, numerous clinical trials have explored the efficacy of mTOR inhibitors in combination with other molecularly targeted or chemotherapeutic agents to reverse drug resistance^47–50^. Besides, while mTOR inhibitors has been reported to suppress CSCs^51^, other studies demonstrated that they are capable of inducing the expansion of drug-resistant CSCs in breast and colorectal cancer ^52,53^. It appears that the effects of mTOR inhibitors on CSC may be context or cell type dependent. Interestingly, induction of ferroptosis is now a novel approach to overcome drug or immunotherapy resistance^54^, as shown by preclinical studies. Therefore, our work highlights that the metabolic status of the cell driven by mTOR pathway modulates the sensitivity to Sal-induced ferroptosis in breast CSCs. Finally, it provides proof-of-concept that careful evaluation of such combination therapy (here co-targeting mTOR inhibition and ferroptosis) is required to develop effective treatments.

## Abbreviations

BCSCs: breast cancer stem cells
CSCs: cancer stem cells
ECAR: extracellular acidification rate
ETC: electron transport chain
FA: fatty acid
FAC: ferric ammonium citrate
FACS: fluorescence activated cell sorting
FTH: ferritin heavy chain
GLS: glutaminase
GO: Gene Ontology
HCQ: hydroxychloroquine
LC-MS: liquid chromatography-mass spectrometry
mt: mitochondrial
mtDNA: mitochondrial DNA
mTOR: the mechanistic target of rapamycin
mTORC1/2: the mechanistic (or mammalian) target of rapamycin complex 1/2
OCR: oxygen consumption rate
OXPHOS: Oxidative Phosphorylation
PUFA: poly-unsaturated fatty acid
RAPTOR: regulatory-associated protein of mTOR
ROS: reactive oxygen species
RT-qPCR: reverse transcription quantitative polymerase chain reaction
Sal: salinomycin
SIN1: mammalian stress-activated protein kinase-interacting protein 1
TCA: tricarboxylic acid
Tf: transferrin
TfR: transferrin receptor.

## DECLARATIONS

### Acknowledgments

The authors gratefully acknowledge funding from INSERM, Université Paris Cité, la ligue nationale contre le cancer, and Comité de Paris de la ligue contre le cancer. We thank Guillaume Andrieu for his feedback on this work, Nicolas Goudin for his help on the mitochondria Analysis and Christine Leroy for her technical assistance.

### Authors’ contributions

Ahmed Hamaï, Maryam Mehrpour, and Emma Cosialls contributed to the study concept and design, acquisition, analysis, interpretation of the data, and manuscript drafting. Emeline Pacreau, Rima Elhage, Clémence Duruel, and Romane Ducloux contributed to data collection. Sylvie Souquere, and Gérard Pierron performed electron microscopy experiments. Sara Ceccacci, Chiara Guerrera performed proteomic experiments and proteomic analysis. Christophe Desterke, and Kevin Roger performed bioinformatic analysis of proteomic data. Ivan Nemazanyy performed metabolomic experiments and metabolite measurements. Mairead Kelly, Elise Dalmas, Yunhua Chang and Vincent Goffin contributed to manuscript drafting. Ahmed Hamaï, Emma Cosialls and Maryam Mehrpour supervised the study.

### Consent for publication

All authors approved the manuscript for submission and consented for publication.

## ADDITIONAL INFORMATION

### Lead contact

Further information and requests for resources and reagents should be directed to and will be fulfilled by the Lead Contact, Dr. Ahmed Hamaï, PhD

### Material availability

All request for resources and reagents should be directed to and will be fulfilled by the lead contact. All reagents will be made available on request after completion of a Materiel Transfer Agreement.

### Data and code availability

All data supporting the findings of this study are available within the paper and are available from the corresponding author upon request.

Any additional information required to reanalyze the data reported in this paper is available from the lead contact upon request.

## SUPPLEMENTARY FIGURES

**Supplementary Fig. S1., related to Figure 1.**
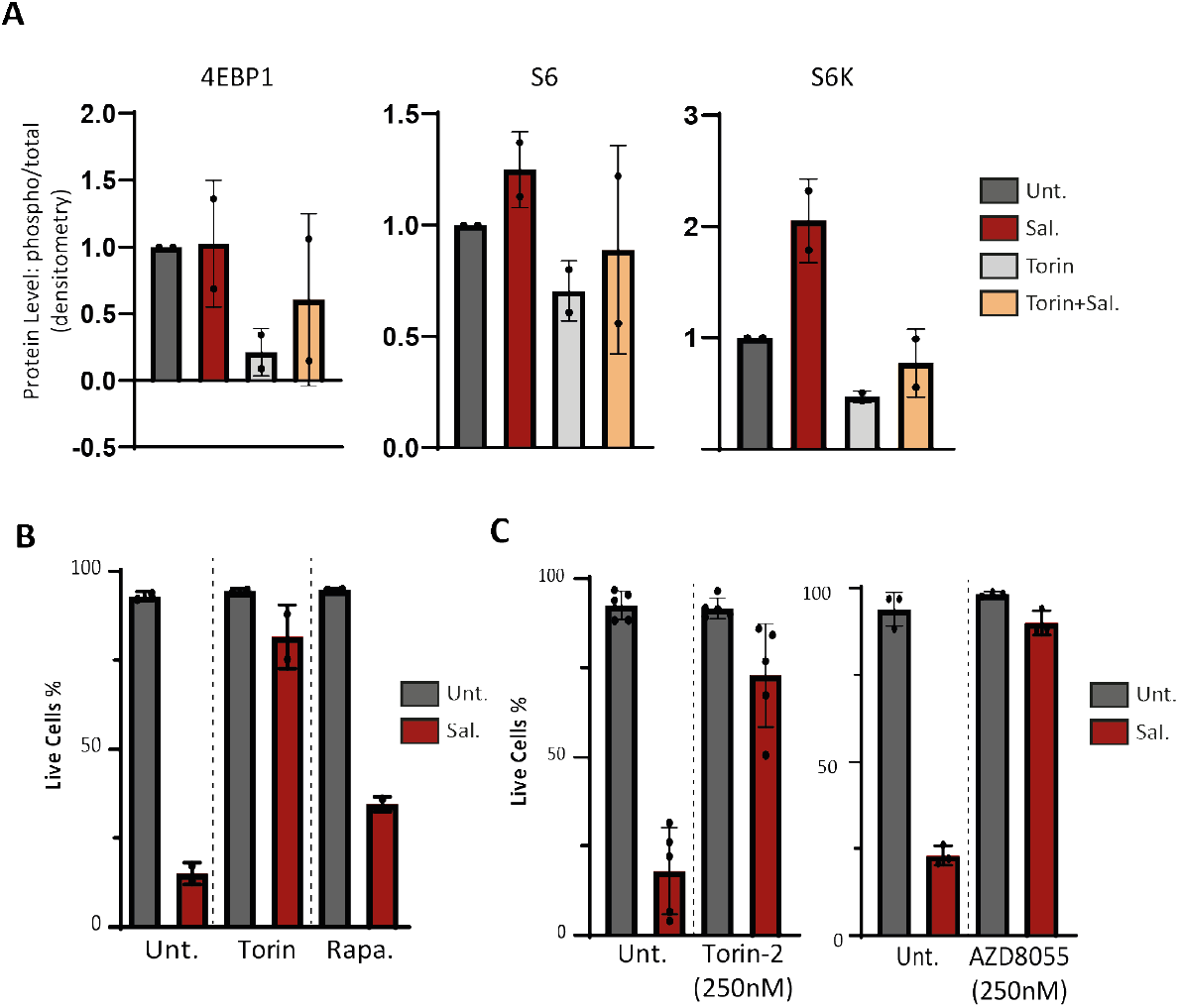
mTOR inhibition inhibits ferroptotic cell death induced by Sal. **(A)** HMLER CD24L were treated with either Sal, Torin or combination of both for 48h. Protein level detected by immunoblot and densitometry analysis normalized on Tubulin level. **(B-C)** HMLER CD24L were treated for 96h with Sal with or without **(B)** Torin 250nM or Rapamycin 250nM **(C)** left panel, Torin-2 (250 nM); right panel, AZD8055 (250 nM). Cell death determined by dapi staining coupled with flow cytometry (FC).

**Supplementary Fig. S2., related to Figure 2.**
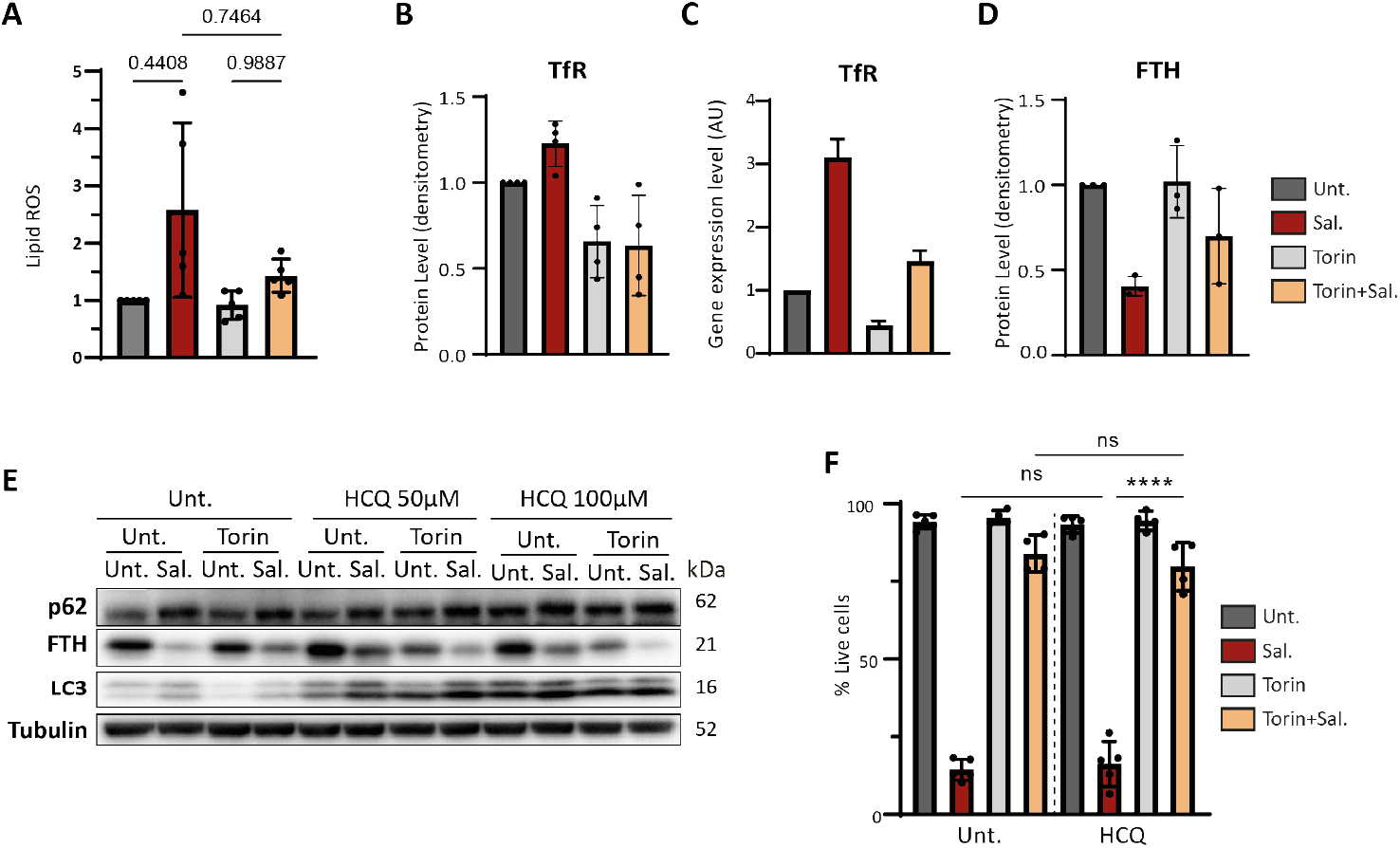
mTOR inhibition impacts iron homeostasis dysregulation induced by Sal. **(A-D)** HMLER CD24L were treated with either Sal, Torin or combination of both for 48h. **(A)** Lipid ROS level determined by BodipyC11 staining coupled with FC (n=5). **(B)** Protein level of TfR detected by immunoblot and densitometry analysis normalized on Tubulin level. **(C)** Gene expression level of TFRC detected by RT-qPCR normalized on Actin level. **(D)** Protein level of FTH detected by immunoblot and densitometry analysis normalized on Tubulin level. **(E)** HMLER CD24L were treated with either Sal or/and Torin in combination with HCQ (50μM or 100 μM) for 48h. Immunoblotting for the indicated autophagy-related protein. Tubulin was used as a loading control. **(F)** HMLER CD24L were treated with either Sal or/and Torin in combination with HCQ (50DDM) for 96h. Cell death determined by dapi staining coupled with flow cytometry (FC). Data are presented: mean ±SD, ANOVA test: ****p<0.0001; ****p<0.002; ****p<0.001

**Supplementary Fig. S3., related to Figure 3.**
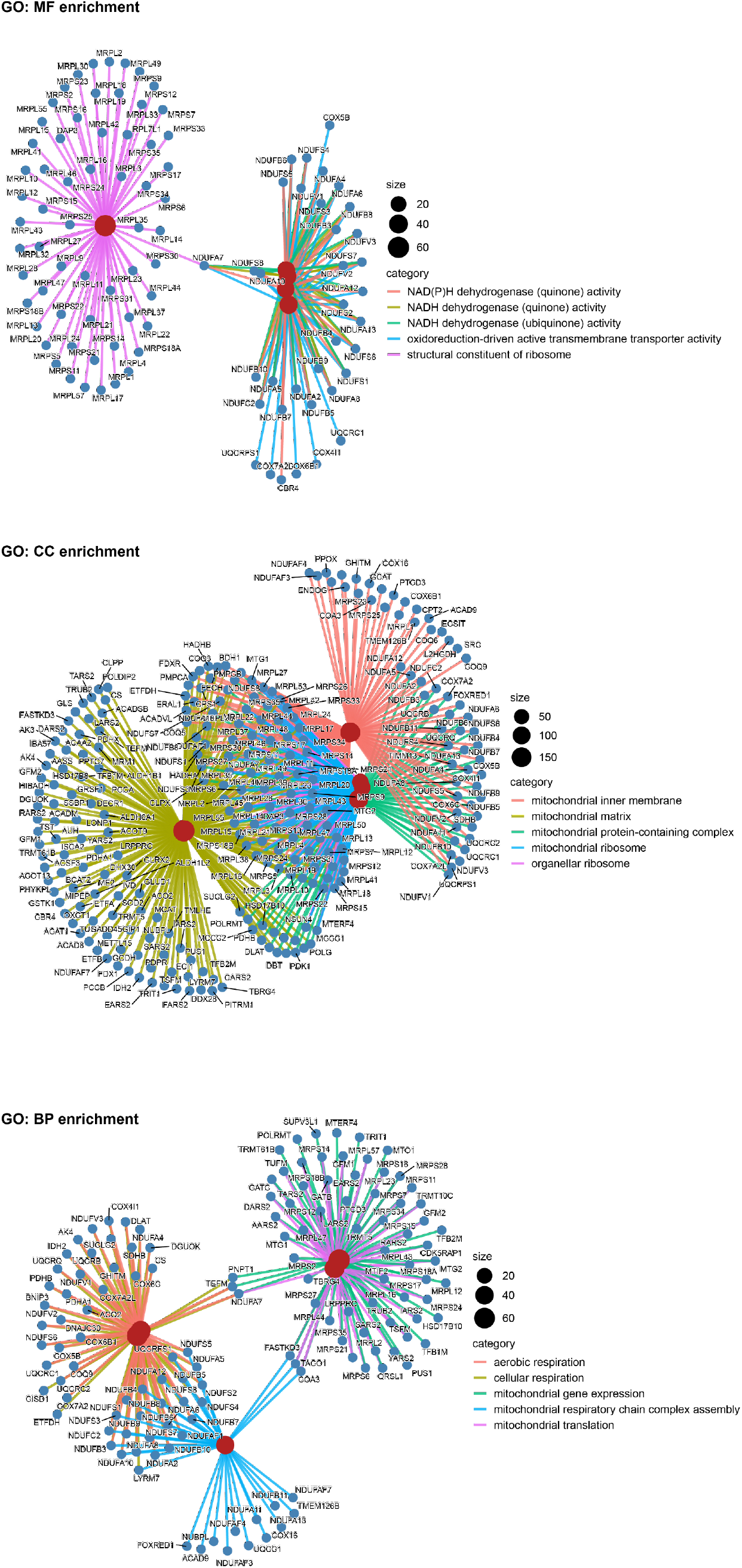
Enrichment map of GO terms enriched in proteins downregulated by Sal treatment (and restored by Sal + Tor combining treatment) from Cluster A shown in an interaction network. MF, Molecular Function; CC, Cellular Component; and BP, Biological Process.

**Supplementary Fig. S4., related to Figure 3.**
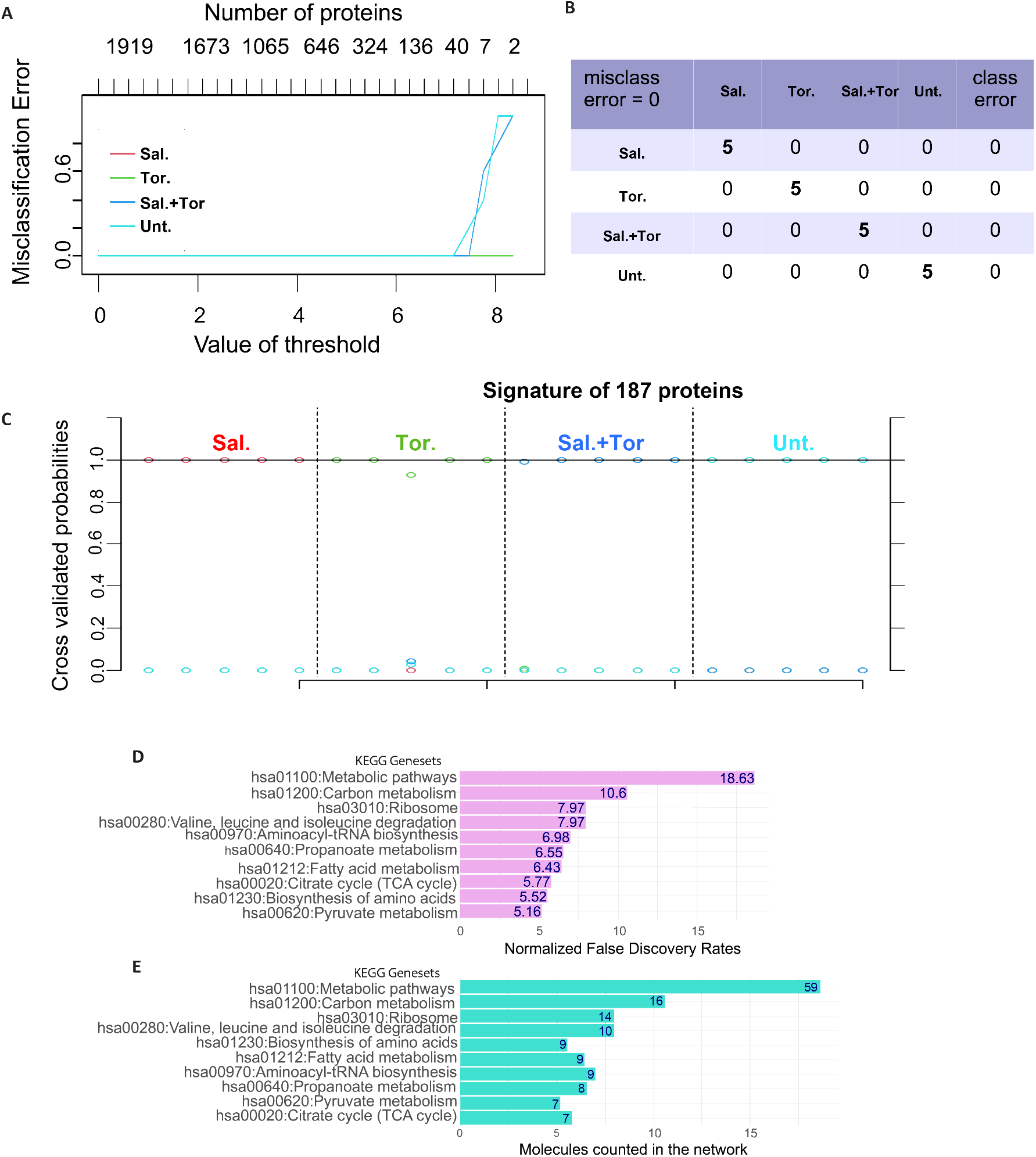
A regulation of 187 proteins discriminates the four experimental conditions investigated by proteomic analysis. **(A)** Supervised machine learning miss-classification error by class according number of proteins included in the model. **(B)** Confusion matrix obtained with the 187-protein signature for optimal threshold of 6. (C) Plot cross validated probabilities for samples obtained with the 187-protein signature. (D) Bar plot of normalized False Discovery Rates obtained on 187 protein-network enriched with KEGG database. (E) Bar plot of molecule counts obtained on 187 protein-network enriched with KEGG database.

**Supplementary Fig. S5., related to Figure 3.**
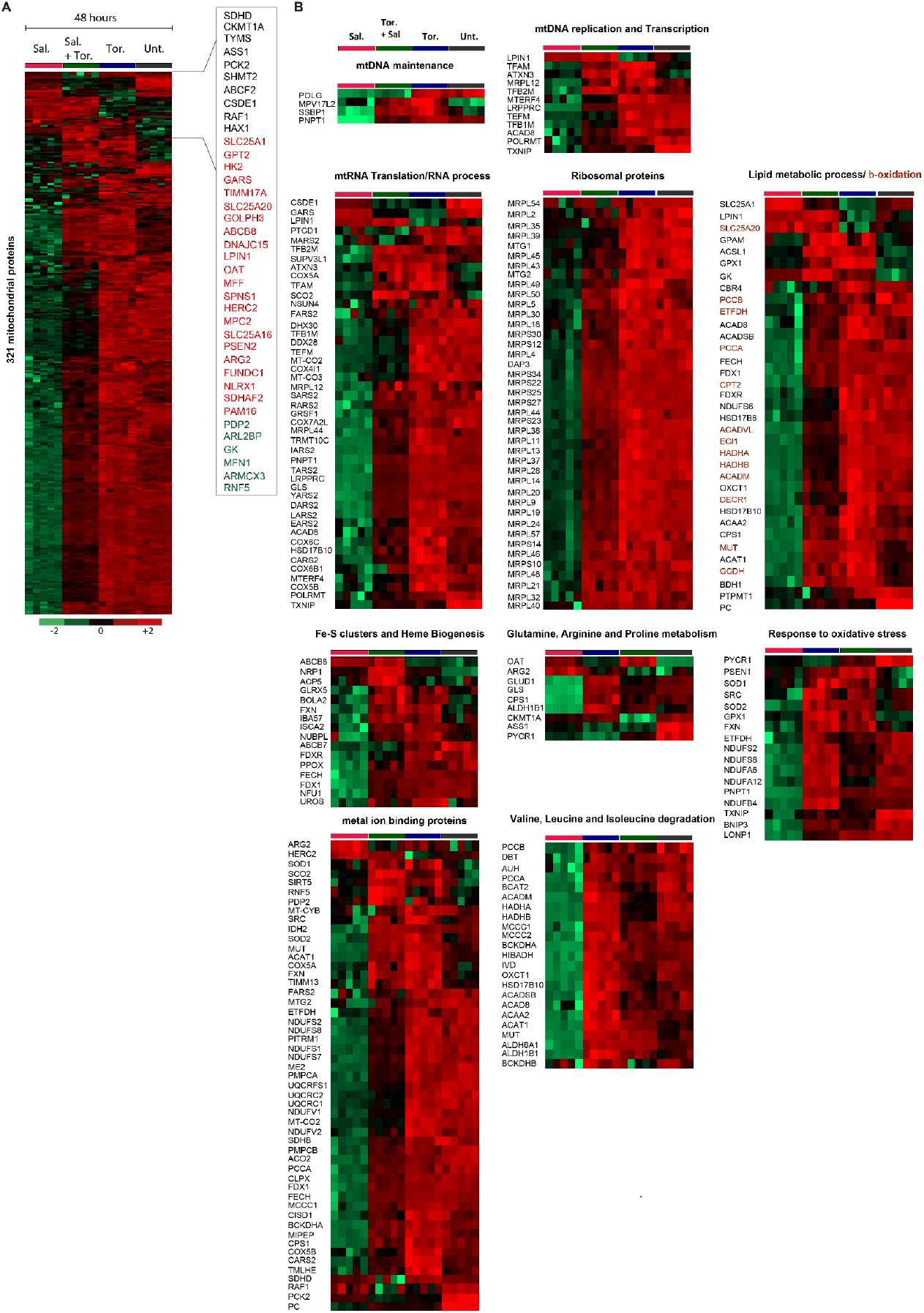
Sal treatment dramatically affects mitochondrial protein expression which is completely rescued by mTOR inhibition. **(A)** Heatmap comparing relative levels of 321 mitochondrial proteins from 1919 proteins significantly modulated by Sal treatment, Tor. alone treatment, and combination treatment compared to untreated (Unt.) cells for 48 h. A focus of some mitochondrial proteins modulated by Sal treatment is shown: red, upregulated; green, downregulated; black, no modulated as compared to untreated cells. **(B)** Heatmap comparing relative levels of proteins from 321 mitochondrial proteins significantly modulated by Sal treatment, Tor. alone treatment, and combination treatment compared to untreated (Unt.) cells for 48 h. These proteins were clustered in function of their role or their category in mitochondria as indicated. Color key indicates protein expression value (green: lowest; red: highest). Proteins were clustered using Perseus software.

**Supplementary Fig. S6, related to Figure 5.**
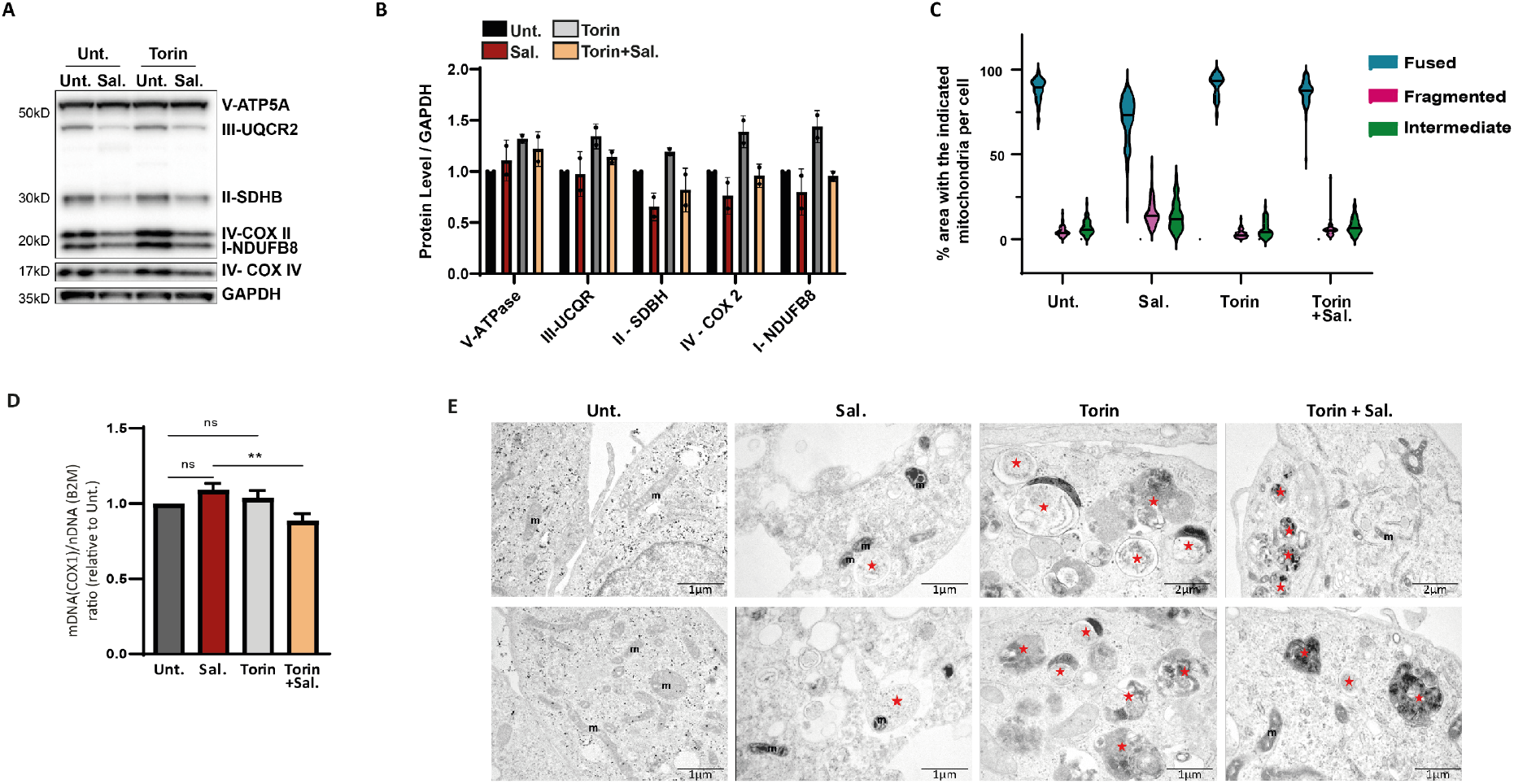
mTOR inhibition protects from mitochondrial damages induced by Sal. **(A-D)** HMLER CD24L were treated with either Sal, Torin or combination of both for 48h. **(A)** Immunoblotting for the indicated ETC-related protein. GAPDH was used as a loading control and **(B)** quantification by densitometry analysis normalized on GAPDH level. **(C)** Quantification of mitochondrial morphology. Analysis on >25 cells from 2 independent experiments. **(D)** The mitochondrial mass was measured by mitoDNA (mDNA, MT-COX1) content. mtDNA was examined by qPCR and normalized by nuclear DNA (nDNA, B2M). Shown are mean ± SEM (n=3 biological replicates). Two tailed and unpaired student’s t test. *, P<0.05, **p < 0.01; ns, not significant. (**E**) TEM images (scale bars: 1μm, 500 nm). Red stars indicate autophagic vacuoles/autolysosomes. m, mitochondria.

## METHODS

### Cell line and Culture

The human mammary epithelial cell line infected with a retrovirus carrying hTERT, SV40, and the oncogenic allele HrasV12, referred to as HMLER CD24L cells, is a subclone known to be rich in the ‘stemness’ phenotype ^55^. HMLER CD24L cells were a generous gift from A. Puisieux (INSERM 1052, Lyon, France). HMLER CD24L cells were cultured in DMEM/F12 + GlutaMAX (Gibco, 31331) supplemented with 10% Fetal Bovine Serum (FBS, Eurobio, CVFSVF00-01), 10 μg/mL Insulin (Sigma, I9278) 0.5 μg/mL hydrocortisone, 10 ng/mL human EGF (Peprotech, AF-100-15), and 0.5 μg/mL puromycin (Invivogen).

### Antibodies, Reagents and Software

| Antibodies (Source, Reference) |
| --- |
| 4EBP1 (Cell Signalling #9644, WB :1/1000) |
| 4EBP1 - phospho (Thr37/46) (Cell Signalling #2855, WB : 1/1000 in 5% BSA) |
| ATG3 (SAB3500500, Sigma Aldrich, WB :1/1000) |
| ATG7 (Sigma Aldrich A2856, WB: 1:1000) |
| ATG9A (PD042, MBL International, WB : 1/1000) |
| BNIP3 (Santa Cruz, sc-56167, WB : 1/1000) |
| BNIP3L (12393, Cell Signaling, WB : 1 :500) |
| DRP1 (BD Biosciences B611113, WB : 1/1000) |
| FTH (Abcam 75973, WB : 1/1000) |
| GAPDH (Santa Cruz sc-47724, WB : 1:2000) |
| Hsp60 (12165, Cell Signaling, WB : 1:500, IF : 1/100) |
| IRP2 (Cell Signalling #37135, WB : 1/1000 in 5% BSA) |
| LC3 (Sigma L7543, WB : 1/1000) |
| MFN2 (Cell Signaling, 9482, WB : 1/1000) |
| NCOA4 (Santa Cruz sc-373739, WB : 1/1000) |
| NDP52 (Proteintech, 12229-1-AP, WB : 1/1000) |
| OPA1 (Cell Signalling, #80471, WB : 1/1000) |
| OPTN (Proteintech 10837-1-AP, WB : 1/1000) |
| p62 (Progen - GP62-C, WB : 1/1000) |
| p70-S6K (Cell signaling #9202, WB : 1/1000) |

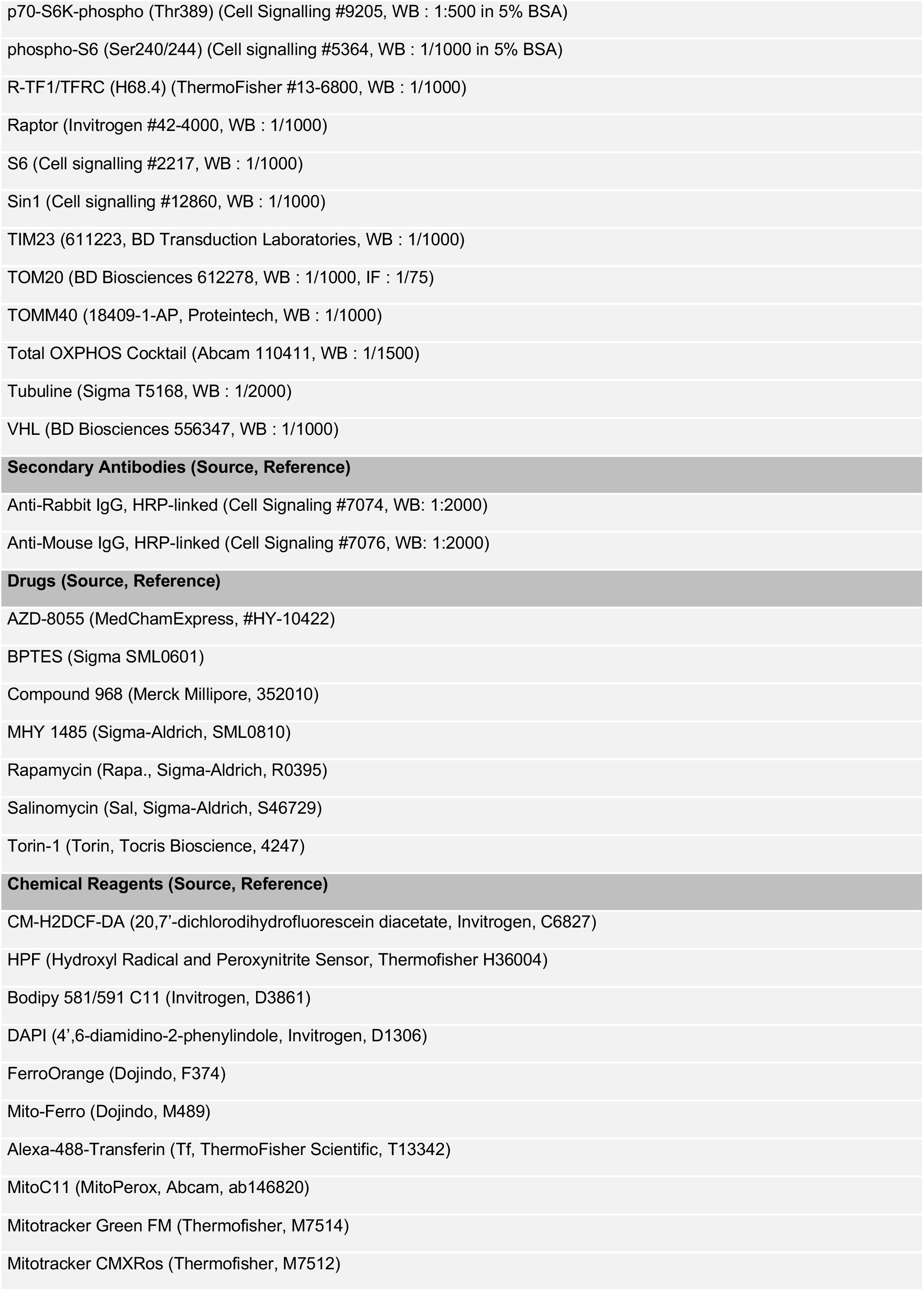

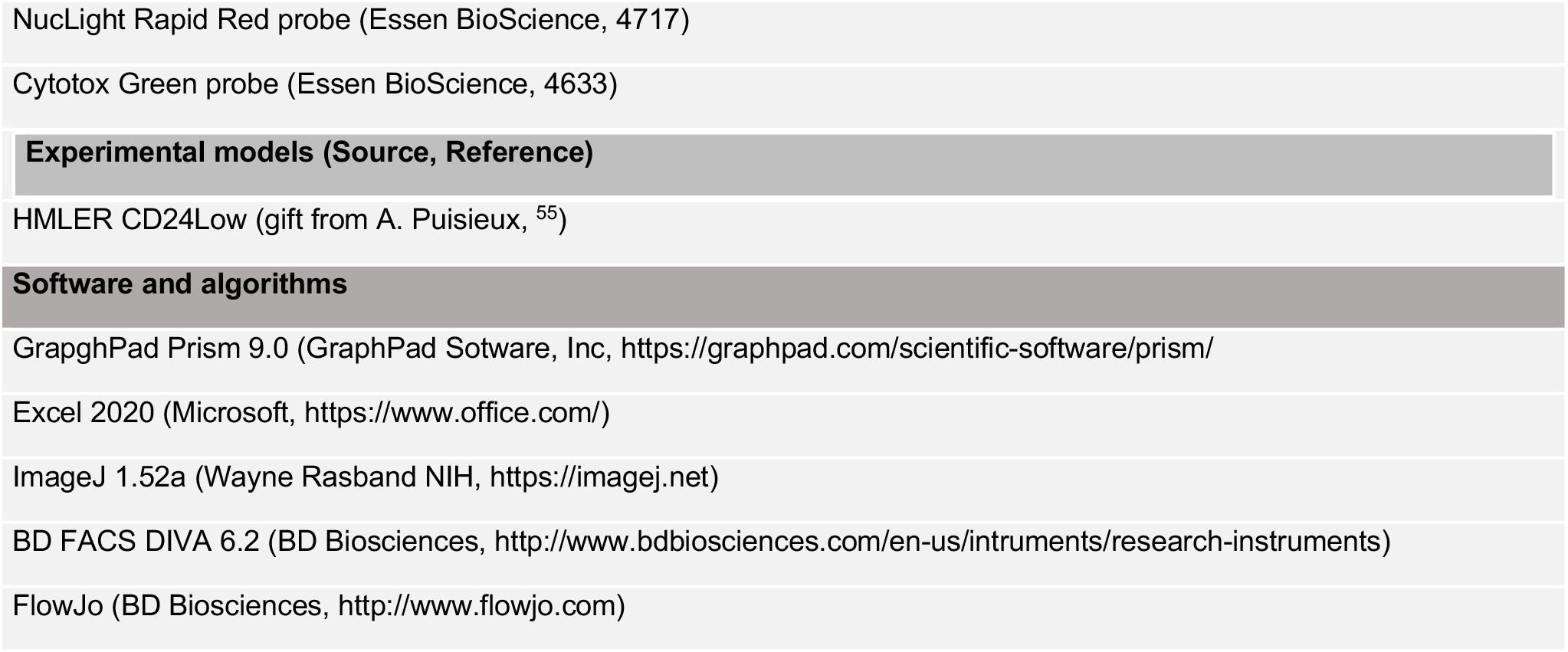

### Drugs Treatments

Unless specified otherwise, cells were seeded at a density of 40,000 cells/well in medium in 12-well plates (or at a density of 1×10^4^ cells/well in 6-well plate for western blot and RT-qPCR). At 70-80% confluence (2-3 days after) cells were treated with the indicated treatments. Drugs are listed in Supplementary Table S3.

### Cell Death (Flow Cytometry)

Cells were treated as indicated during the desired time, then detach with trypsin, collected and rinse with PBS. Subsequently, DAPI (1:2000 in PBS) was added for dead cell quantification using a flow cytometer (Fortessa, BD Biosciences). A minimum of 50,000 cells was analyzed per condition.

### Cell Death (IncuCyte)

Cells were seeded in 24-well plates at a density of 20,000 cells by well and treated with the indicated treatments. Simultaneously, NucLight Rapid Red probe (4717, Essen BioScience) and Cytotox Green probe (4633, Essen BioScience) were added. Images (20X) of the same fields were taken every 2 h for 96 h by a real-time IncuCyteS3 Live-Cell analysis system (Essen Bioscience, Ann Arbor, Michigan, USA). Fluorescence intensities were analyzed using the IncuCyte software, and results were displayed normalized to the initial time point (time t=0).

### Measurement of Fluorescent probes staining

Cells were treated as indicated during the desired time, then detach with trypsin, collected and rinse with PBS. Subsequently, probes diluted in HBSS were added (with the indicated concentration) and incubated during 30mn (except if specified) at 37°C + 5% CO_2_. Excess probe was removed by washing the cells with PBS, and DAPI (1:2000) was added to stain dead cells. Fluorescence signals were measured using a flow cytometer (Fortessa, BD Biosciences). A minimum of 50,000 cells were analyzed for each condition. Data were processed using BD FACSDiva software and analysed using FlowJO software on median fluorescence level gated on live cells. The probes (with their concentration and their incubation time if different from 30mn) are listed in Supplementary Table S3.

### Proteomics

#### Sample Preparation for Proteomic Analysis

S-TrapTM micro spin column (Protifi, Hutington, USA) digestion was performed on 20 µg of cell lysates according to manufacturer’s instructions and optimized as described in Ceccacci et al., 2022 ^56^. After elution, peptides were vacuum dried and resuspended in 50 µl of 2 % ACN, 0.1 % FA and quantified by Nanodrop ^56^.

#### nanoLC-MS/MS Protein Identification and Quantification

400 ng of each sample was injected on a nanoElute (Bruker Daltonics, Germany) HPLC (high-performance liquid chromatography) system coupled to a timsTOF Pro (Bruker Daltonics, Germany) mass spectrometer. HPLC separation (Solvent A: 0.1% formic acid in water; Solvent B: 0.1% formic acid in acetonitrile) was carried out at 250 nL/min using a packed emitter column (C18, 25 cm×75μm 1.6μm) (Ion Optics, Australia) using a gradient elution (2 to 13% solvent B during 41min; 13 to 20% during 23min; 20% to 30% during 5min; 30% to 85% for 5min and finally 85% for 5min to wash the column). Mass-spectrometric data were acquired using the parallel accumulation serial fragmentation (PASEF) acquisition method. The measurements were carried out over the m/z range from 100 to 1700 Th. The range of ion mobilities values from 0.75 to 1.25 V s/cm^2^(1/k0). The total cycle time was set to 1.17s and the number of PASEF MS/MS scans was set to 10.

#### MS Data Processing and Bioinformatics Analysis

The obtained data were analyzed using MaxQuant version 2.0.1.0 and searched with Andromeda search engine against the UniProtKB/Swiss-Prot *Homo sapiens* database (release 02-2021, 20408 entries). To search parent mass and fragment ions, we set a mass deviation of 3 ppm and 20 ppm respectively. The minimum peptide length was set to 7 amino acids and strict specificity for trypsin cleavage was required, allowing up to two missed cleavage sites. Carbamidomethylation (Cys) was set as fixed modification, whereas oxidation (Met) and N-term acetylation were set as variable modifications. The false discovery rates (FDRs) at the protein and peptide level were set to 1%. Scores were calculated in MaxQuant as described previously ^57^. The reverse and common contaminants hits were removed from MaxQuant output. Proteins were quantified according to the MaxQuant label-free algorithm using LFQ intensities; protein quantification was obtained using at least 1 peptide per protein. Match between runs was allowed.

Statistical and bioinformatic analysis, including heatmaps, profile plots and clustering, were performed with Perseus software (version 1.6.15.0) freely available at ^58^. Intensities were log2 transformed for statistical analysis. For statistical comparison, we set 4 groups, each containing 5 biological replicates. We then filtered the data to keep only proteins with at least 5 valid values in at least one group. Next, the data were imputed to fill missing data points by creating a Gaussian distribution of random numbers with a standard deviation of 33% relative to the standard deviation of the measured values and 1.8 standard deviation downshift of the mean to simulate the distribution of low signal values. We performed an ANOVA test, FDR<0.01, S0=1. Hierarchical clustering of proteins that survived the test was performed in Perseus on logarithmised LFQ intensities after Z-score normalization of the data, using Euclidean distances.

### Targeted liquid chromatography mass spectrometry analysis of metabolites

For targeted metabolomics analysis, 2 × 10^5^ WT and KD.7 cells were treated or untreated with 100 ng/mL OSM for 24, 48, and 72 h. Each sample was washed three times with cold DPBS, frozen in liquid nitrogen, and stored at -80 °C. Metabolites were extracted with an adequate volume (calculated from cell count 2 × 10^6^ cells/mL) of an aqueous solution of methanol and acetonitrile (20:50:30). Samples were vortexed for 5 min at 4 °C and then centrifuged at 16,000 g for 15 min at 4 °C. The supernatants were collected and stored at -80 °C until analyses. LC/MS analyses were conducted on a Q Exactive Plus Orbitrap mass spectrometer equipped with an Ion Max source and a HESI II probe and coupled to a Dionex UltiMate 3000 UPLC system (Thermo Fischer Scientific). External mass calibration was performed as recommended by the manufacturer. A 5 µL aliquot of each sample was injected onto a ZIC-pHILIC column (150 mm × 2.1 mm i.d. 5 μm, Millipore) fitted with a guard column (20 mm × 2.1 mm i.d. 5 μm, Millipore) for the liquid chromatography separation. The chromatographic gradient was run at a flow rate of 0.200 μl/min with Buffer A (20 mM ammonium carbonate and 0.1% ammonium hydroxide, pH 9.2) and Buffer B (acetonitrile) as follows: at 0–20 min, a linear gradient from 80% to 20% B was administered; at 20–20.5 min, a linear gradient from 20% to 80% B was administered; at 20.5–28 min, 80% B was maintained. The mass spectrometer was operated in full-scan, polarity-switching mode, with the spray voltage set to 2.5 kV and the heated capillary held at 320 °C. The sheath gas flow was set to 20 units, the auxiliary gas flow was set to 5 units, and the sweep gas flow was set to 0 units. The metabolites were detected across a mass range of 75–1000 m/z at a resolution of 35,000 (at 200 m/z), with the AGC target at 10^6^ and the maximum injection time at 250 ms. Lock masses were used to ensure a mass accuracy below 5 ppm. Data were acquired with ThermoXcalibur software. The peak areas of the metabolites were determined using Thermo Trace Finder software and identified by the exact mass of each singly charged ion and by the known retention time on the HPLC column. Statistical and pathway analyses were performed using Metaboanalyst 5.0 software.

### Measurement of OCR and ECAR

OCR and ECAR were analyzed using the Seahorse XFe96 bioenergetic analyzer in accordance with the manufacturer’s instructions (Agilent Technologies). Briefly, HMLER CD24^low^ were seeded at a density of 10^4^ cells per well in a specialized 96-well Seahorse XF96 V3 PS microplate (101085-004, Agilent Technologies). 48h after, cells were incubated with 500nM Salinomyin and/or 250nM of Torin. Cells were incubated for 1 h in unbuffered XF assay media (Agilent Technologies) supplemented sequentially with either 2 mM glutamine, 25 mM glucose (G8769, Sigma Aldrich), and 1 mM sodium pyruvate for OCR analysis. For OCR/ECAR measurements, the XF Cell Mito Stress Test (103015-100, Agilent Technologies) was used. Compounds were injected during the assay at the following final concentrations: oligomycin (75351, Sigma Aldrich, ATP synthase inhibitor): 1 µM; FCCP (370-86-5, Sigma Aldrich, uncoupling agent measuring the maximal respiration capacity): 0.5 µM; antimycin A (A8674, Sigma Aldrich, mETC inhibitor): 1 µM.

### Small interfering RNA transfection

HMLER CD24L cells were seeded at a density of 30 000 cells per well in a 12-well plate. After 24h of pre-incubation cells were transfected at sub-confluence with 10nM of si-CTRL (D001810-10-20, ThermoFisherScientific, ON-TARGET™*plus* control)), si-RAPTOR (sc-44069, Santa Cruz), si-SIN1 (sc-60984, Santa Cruz) were mixed with 3μL/well of Lipofectamine RNAiMAX (13778-150, Invitrogen) diluted in Opti-MEM (51985-042, Gibco) following the manufacturer’s instructions. After 4h of transfection, mix was replaced with fresh medium. The next day a second transfection with the same siRNAs was performed using the same protocol, but after 4h of transfection, 500uL of fresh medium was added. Cells were treating cells the next day as indicated in Figure 1. Cells were then collected and analyzed by flow cytometry and/or Western blot at corresponding time.

### RNA analysis

Total RNA was extracted from cell pellets using the Nucleospin RNA kit (740955, Macherey-Nagel - Hoerdt) according to the manufacturer’s protocol and quantified using a NanoDrop 2000 spectrophotometer (Thermo Fisher Scientific). cDNA was generated from the total RNA (250 ng) using random hexamer primers (S0142, Thermo Fisher Scientific) and the M-MLV reverse transcriptase (28025-016, Invitrogen) according to the manufacturer’s protocol. mRNA levels of target genes were quantified by qPCR using iTaq Universal SYBR Green Supermix (172-5124, BioRad) according to the manufacturer’s protocol in a CFX96 thermal cycler (BioRad). The data were normalized to the internal control β-actin. Relative gene expression levels were calculated using the 2-ΔΔCt method. Primers used: FTH1 (Forward: 5’-CTGGAGCTCTACGCCTCCTA-3’; Reverse: 5’-TCTCAGCATGTTCCCTCTCC-3’); NCOA4 (Forward: 5’-CAGCTGGTGAGTCGGTGAC-3’; Reverse: 5’-TCCGTGCATCACTACACCTC-3’); TfR-1 (Forward: 5’-GAGCCTGTGGGGAAGGG-3’; Reverse: 5’-AGGCTGAACCGGGTATATGA-3’); β-actin (Forward: 5’-AAGACCTGTACGCCAACACA-3’; Reverse: 5’-TGATCTCCTTCTGCATCCTG-3’).

### Immunoblotting

Cell lysates were prepared on ice in an appropriate lysis buffer (50 mM Tris-HCl, pH 7.5, 150 mM NaCl, 1% TRITON X-100, 0.5% NP-40, 10% glycerol, 1% protease, and a Phosphatase Inhibitor Cocktail (78442, Thermo Fisher Scientific)). Protein concentrations were determined with the Pierce BCA protein assay kit (23225, Thermo Fisher Scientific). Equal protein amounts (15-20 µg) diluted in a 5x Laemmli buffer were denatured by heating at 95 °C for 5 min and separated by electrophoresis on 4%-12% NuPAGE Bis-Tris Gel, and then transferred onto a 0.45 μm a PVDF membrane. Membranes were blocked with 5% non-fat dry milk in PBS-T (D-PBS with 0.1% Tween-20) for 1h at room temperature, and then incubated with primary antibodies at 4°C overnight. Membranes were then washed with DPBS-T, and incubated with the appropriate HRP-coupled secondary antibody for 1h30 at RT. Antibodies are listed in Supplementary Table S3. Membranes were then washed with PBS-T, and bound antibodies were detected using an ECL detection kit (Immobilon Western ECL, Millipore) or with the ChemiDoc Imaging Systems (BioRad) using the CCD camera for light capture according to the manufacturer’s protocol. Signals were quantified using Image Lab Software (Bio-Rad) and normalized to Tubulin or GAPDH.

### Transmission electron microscopy

Cells were treated in 6-well plate as indicated during 48h. For ultrastructural analysis, cells were fixed with 1.6% glutaraldehyde in 0.1 mol/L phosphate buffer. After scraping, cell pellets were secondary fixed with 2% osmium tetroxide and dehydrated using ethanol. Cells were embedded in Epon 812 resin. Polymerization was complete after 48 h at 60°C. Ultrathin sections were collected on 100-mesh grids coated with Formvar and carbon, stained with standard uranyl acetate and lead citrate solutions. The sections were then examined under a FEI Technai Spirit transmission electron microscope at 80 Kv, and images were acquired with a SIS Mega view III CCD camera.

### Immunofluorescence microscopy

Cells were seeded at a density of 25,000 cells per well on glass coverslips in a 12-well plate, at sub-confluence cell were treated. Then after 48h, cells were washed twice with PBS and fixed with 4% paraformaldehyde (15714, Electron Microscopy Sciences) for 13 min at 37°C. After washing twice with PBS, cells were permeabilized with PBS+10% FBS and 0.3% TRITON X-100 (X100, Sigma Aldrich) at room temperature for 20 min. After washing twice with a washing buffer (PBS + 10% FBS), cells were incubated at 4 °C overnight with the indicated primary antibodies, including anti-TOM20 (1:75), anti-Hsp60 (1:100). Cells were then washed three times and incubated with Alexa 488/647-labeled anti-mouse/rabbit secondary antibody (Invitrogen, A21202, A31573) for 1h at RT. All antibodies were diluted with washing buffer. The cells were then washed twice, incubated with DAPI (1:2000) 10mn in PBS, wash twice with PBS and slides were mounted with a coverslip using fluoromount-G medium (0100-20, SouthernBiotech). After adhering, slides were allowed to dry overnight and stored at 4 °C in dark to prevent photobleaching. Cell images were obtained using a Spinning Disk Zeis or a Confocal Leica SP8 with objective 63X oil. Stacks of images were collected every 0.22um (or 0.01 μm for the TOM20) along the z-axis. Images were processed by ImageJ software.

### Mitochondria Analysis

Mitochondrial morphology qualification and quantification was performed using two supervised machine-learning (ML) modules to segmented mitochondria and to classified them on their morphology. Firstly, images were prepared with a FIJI macro (citation). Then, with the “pixel classification + object classification” ML module of Ilastik (V1.3.3post3 citation), on 3D images of z-stack of TOM20, mitochondrial network was segmented into objects. The segmentation was carried out using a supervised ML trained to identify three classes of object: background, mitochondria edges and mitochondria body, based on several properties/features: Color/Intensity, Edge, Texture. The ML was performed for each experiment on at least 10 cells by condition, before being applied to each experiment independently. Mitochondria were then segmented using a Hysterisis thresholding approach. The mitochondria body recognized as labelled object were saved as a new 3D images. Then, we used Imaris V9.9 Software (Oxford Instruments). Following, mitochondria body were classified into three classes: fragmented, hyperfused and intermediate, using a supervised ML training based on several shapes features. This ML was performed on at least 20 cells/condition, before being applied to all the experiments. Finally for each cell, total volume of mitochondria was compared with volume of each class. Data were analysed with Graphpad Prism and showed by cell the proportion of the area for each class.

### Mitochondrial DNA quantification

Cells were treated as indicated during the desired time, then detach with trypsin, collected and rinse with PBS. The total DNA was extracted by using DNeasy Blood & Tissue Kit (# 69,504, Qiagen, Germany), following manufacturer’s instructions. Quantification of mitochondrial DNA (mtDNA) and nuclear DNA (nDNA) were performed by qPCR with SYBR green-based detection (Thermo Fisher Scientific) using iTaq Universal SYBR Green Supermix (172-5124, BioRad) according to the manufacturer’s protocol in a qTOWER 2.0/2.2 thermal cycler (Analytic Jena). Relative mDNA:nDNA ratio was calculated using the 2^-ΔΔCt^ method upon targeting of nuclear-encoded gene (human B2M) and mitochondrial-encoded gene (human COX1) (Quiros, PM., Goyal A. et al., Curr. Protoc. Mouse Biol.). The sequences of the primers are the following: forward primer: 5’-CCCACCGGCGTCAAAGTAT -3’ and reverse primer: 5’-TGCAGCAGATCATTTCATATTGC -3’ for COX1; and forward primer: 5’-TGCTGTCTCCATGTTTGATGTATCT -3’ and reverse primer: 5’-TCTCTGCTCCCCACCTCTAAGT -3’ for B2M.

### Bioinformatics analysis on proteomic data

Bioinformatics analyses were performed in R software environment version 4.2.1. Based on annotated proteomic matrix and taking in account the experimental groups (Unt.: Untreated; Torin: Tor.; Salinomycin: Sal; Torin + Salinomycin: TorSal) a supervised leave one out process of machine learning was performed by shrunken centroid determination with pamr R-package ^59^ version 1.56.1. Minimal signature with optimal threshold was determined for a minimal missclassication error of the samples between supervised experimental groups. A quick decrease of misclassification error was observed in four experimental groups (**Supplementary Fig. S4A**). A minimal proteomic signature with threshold fixed to 6 of value (**Supplementary Fig. S4A**) was retained. Minimal signature was validated by unsupervised principal component analysis performed with « prcomp » r-base function and plotted with « autoplot » function from ggfortify R-package version 0.4.14. This minimal proteomic signature allowed to obtain a perfect classification of the groups after cross validation by leave one out process (**Supplementary Fig. S4B**). Based on this signature, a good stratification of the samples between groups according their cross-validated probabilities was performed (**Supplementary Fig. S4C**). Minimal signature was also validated by unsupervised clustering (Pearson distances) performed with heatmap R-package version 1.0.12. Network functional enrichment was done with Kyoto Encyclopedia of Genes and Genomes (KEGG) database ^60^.

### Quantification and statistical analysis

All results were expressed as mean values D SD and were compared by one-way or two-way ANOVA or an unpaired two-tailed Student’s t-test. Analyses were performed using Graph Pad Prism 9.0. Differences were considered to be statistically significant when P<0.05.

